# Loss of Piccolo function in rats induces Pontocerebellar Hypoplasia type 3-like phenotypes

**DOI:** 10.1101/774422

**Authors:** Joanne Falck, Christine Bruns, Sheila Hoffmann, Isabelle Straub, Erik J. Plautz, Marta Orlando, Humaira Munawar, Marion Rivalan, York Winter, Zsuzsanna Izsvák, Dietmar Schmitz, F. Kent Hamra, Stefan Hallermann, Craig Garner, Frauke Ackermann

## Abstract

Piccolo, a presynaptic active zone protein, is best known for its role in the regulated assembly and function of vertebrate synapses. Genetic studies suggest a further link to several psychiatric disorders as well as Pontocerebellar Hypoplasia type 3 (PCH3), although a causal relationship is lacking. We have characterized recently generated knockout (*Pclo^gt/gt^*) rats. Analysis revealed a dramatic reduction in brain size compared to wildtype (*Pclo^wt/wt^*) animals, attributed to a decrease in the size of the cerebral cortical, cerebellar and pontine regions. Analysis of the cerebellum and brainstem revealed a reduced granule cell (GC) layer and a reduction in size of pontine nuclei. Moreover, the maturation of mossy fiber (MF) afferents from pontine neurons and the expression of the α6 GABA_A_ receptor subunit at the MF-GC synapse are perturbed, as well as the innervation of Purkinje cells by cerebellar climbing fibers (CFs). Ultrastructural and functional studies revealed a reduced size of MF boutons, with fewer synaptic vesicles and altered synaptic transmission. These data imply that Piccolo is required for the normal development, maturation and function of neuronal networks formed between the brainstem and cerebellum. Consistently, behavioral studies demonstrated that adult *Pclo^gt/gt^* rats display impaired motor coordination, despite adequate performance in tasks that reflect muscle strength and locomotion. Together these data suggest that loss of Piccolo function in patients with PCH3 could be causal for many of the observed anatomical and behavioral symptoms, and that the further analysis of these animals could provide fundamental mechanistic insights into this devastating disorder.

**Significance Statement:** Pontocerebellar Hypoplasia type 3 is a devastating developmental disorder associated with severe developmental delay, progressive microcephaly with brachycephaly, optic atrophy, seizures and hypertonia with hyperreflexia. Recent genetic studies have identified non-sense mutations in the coding region of the Piccolo gene, suggesting a functional link between this disorder and the presynaptic active zone. Our analysis of Piccolo knockout rats supports this hypothesis, formally demonstrating that anatomical and behavioral phenotypes seen in patients with PCH3 are also exhibited by these Piccolo deficient animals.

## Introduction

Pontocerebellar Hypoplasia is a rare and highly heterogeneous group of neurological disorders, often with a genetic origin, characterized by an abnormally small cerebellum and pons (Rajab et al., 2003; Namavar et al., 2011). In type 3 Pontocerebellar Hypoplasia (PCH3) - also known as Cerebellar Atrophy with Progressive Microcephaly (CLAM) - patients suffer from severe developmental delay, progressive microcephaly with brachycephaly, seizures, hypertonia with hyperreflexia and short stature (Rajab et al., 2003; Namavar et al., 2011). Additional features include the presence of craniofacial dysmorphisms and optic atrophy (Durmaz et al., 2009; Rudnik-Schoneborn et al., 2014).

Previous studies mapped the PCH phenotype to chromosome 7q11-21 (Rajab et al., 2003; Durmaz et al., 2009). More recently, a single nucleotide polymorphism (SNP) in the human *Pclo* gene - indeed, located on chromosome 7 at position 21.11 - has been found in patients with PCH3. This non-sense mutation is predicted to eliminate the C-terminus of the longest Piccolo isoforms including its PDZ and C2 domains (Ahmed et al., 2015) and perhaps destabilize the protein, leading to the hypothesis that Piccolo loss of function is responsible for the phenotypes seen in this neurodevelopmental/neurodegenerative disorder.

Piccolo is a very large (560kDa) multidomain presynaptic scaffold protein and core component of the cytoskeletal matrix assembled at active zones (CAZ) (Cases-Langhoff et al., 1996). A range of studies suggest that Piccolo uses its multidomain structure to scaffold not only other CAZ proteins critical for the regulated release of neurotransmitters, but also proteins involved in the dynamic assembly of F-actin, synaptic vesicle (SV) recycling and synapse integrity (Gundelfinger et al., 2015; Ackermann et al., 2019). Intriguingly, Piccolo is present at nearly every synaptic subtype including glutamatergic, GABAergic, cholinergic and dopaminergic synapses within the central (CNS) and peripheral nervous system (PNS) (Cases-Langhoff et al., 1996; Fenster et al., 2000; Fenster and Garner, 2002) and is highly expressed in the cerebrum, hippocampus, cerebellum and olfactory bulb, among others (Cases-Langhoff et al., 1996; Human Protein Atlas, 2015). It is one of the very first active zone (AZ) proteins recruited to nascent synapses *in vitro* as well as in the developing brain (Zhai et al., 2001). For example, Piccolo appears at emerging synapses formed between mossy fiber boutons and cerebellar granule cells as well as between parallel fiber boutons and Purkinje cell dendrites during the earliest stages of cerebellar development (Zhai et al., 2001). The large size of Piccolo and the complexity of the *Pclo* gene has thwarted most efforts to elucidate its function, though critical roles in retinal ribbon synapse formation and visual function (Regus-Leidig et al., 2014; Muller et al., 2019) as well as the integrity of hippocampal synapses has been identified (Waites et al., 2013). What remains unclear is how Piccolo contributes to cerebellar development and whether, as suggested by genetic studies, it has a primary role in the etiology of PCH3. The recent generation of a Piccolo knockout rat (*Pclo^gt/gt^*) using transposon mutagenesis (Medrano et al., 2019) provides an opportunity to explore this potential relationship.

In the current study, we have assessed the contribution of Piccolo to cerebellar structure and function through the anatomical, functional and behavioral characterization of adult *Pclo^gt/gt^* rats. Our analysis reveals a striking number of similarities to patients with PCH3, including a smaller cerebral cortex, a reduced volume of the cerebellum and pons as well as impaired motor control and the presence of seizures. Our analysis has also uncovered changes in the anatomical and ultrastructural features of mossy fiber terminals and electrophysiological properties of these synapses. Together, these phenotypes are predicted to not only alter the functionality of the cerebellum but to contribute to motor and perhaps also behavioral dysfunctions seen in PCH3 affected individuals.

## Results

### Brain morphology is changed in Piccolo knockout brains

The recent analysis of a pair of boys with PCH3 identified a non-sense mutation in the coding region of the human *Pclo* gene (chr7:82579280 G>A), predicted to eliminate the C-terminal third of the longest Piccolo isoforms (Ahmed et al., 2015) and likely its expression. These individuals have profound cognitive and motor impairment as well as atrophy of the cerebrum, cerebellum and pons (Ahmed et al., 2015). A fundamental question is whether Piccolo loss of function is causal for this disorder. To explore this possibility, we have characterized a recently generated line of rats (*Pclo^gt/gt^*) wherein transposon mutagenesis was used to disrupt the Piccolo gene via an insertion into exon 3 (Figure 1A). This insertion is predicted to cause a frame shift in the reading frame and thus disrupt the expression of full-length Piccolo (560kDa) and most of its alternatively spliced lower molecular weight isoforms (70-350kDa). Schematic is adapted from Ackermann et al. (Ackermann et al., 2019; Medrano et al., 2019). Just after birth (P0-P2) Piccolo pups were found born in normal Mendelian numbers (Figure 1B). Western blot analysis of brain lysates from postnatal day 0-2 (P0-P2) pups demonstrates the loss of nearly all isoforms in homozygous knockout animals (Figure 1C). To assess whether Piccolo loss of function adversely affects brain development, we performed an anatomical characterization of *Pclo^wt/wt^* and *Pclo^gt/gt^* animals; *Pclo^gt/gt^* pups were smaller and weighed significantly less than their *Pclo^wt/wt^* littermates (Figure 1D, E and F). However, brain weights were not significantly altered between *Pclo^gt/gt^* and *Pclo^wt/wt^* littermates at P0-2 and display similar brain morphology (Figure 1G, H and I), suggesting that changes in brain size develop postnatally. However, the overall size and weight of *Pclo^gt/gt^* brains was significantly reduced in 3 month-old adult rats compared to *Pclo^wt/wt^* brains (Figure 1J and K). To assess whether this was associated with an overall loss in brain volume or due to reductions in specific brain regions, serial sagittal and coronal sections from 3 month-old animals were collected and stained with Nissl to visualize brain morphology (Figure 1L). Qualitative analysis revealed that thalamic, cerebellar and brainstem regions are dramatically reduced in size in 3 month-old *Pclo^gt/gt^* animals. Some thinning of the cerebral cortex can also be observed, however, no obvious changes in the hippocampus are detectable. Furthermore, ventricles (V1, V2, V3 and V4) in *Pclo^gt/gt^* brains were all notably larger than in *Pclo^wt/wt^* littermates (Figure 1L).

**Figure 1.**
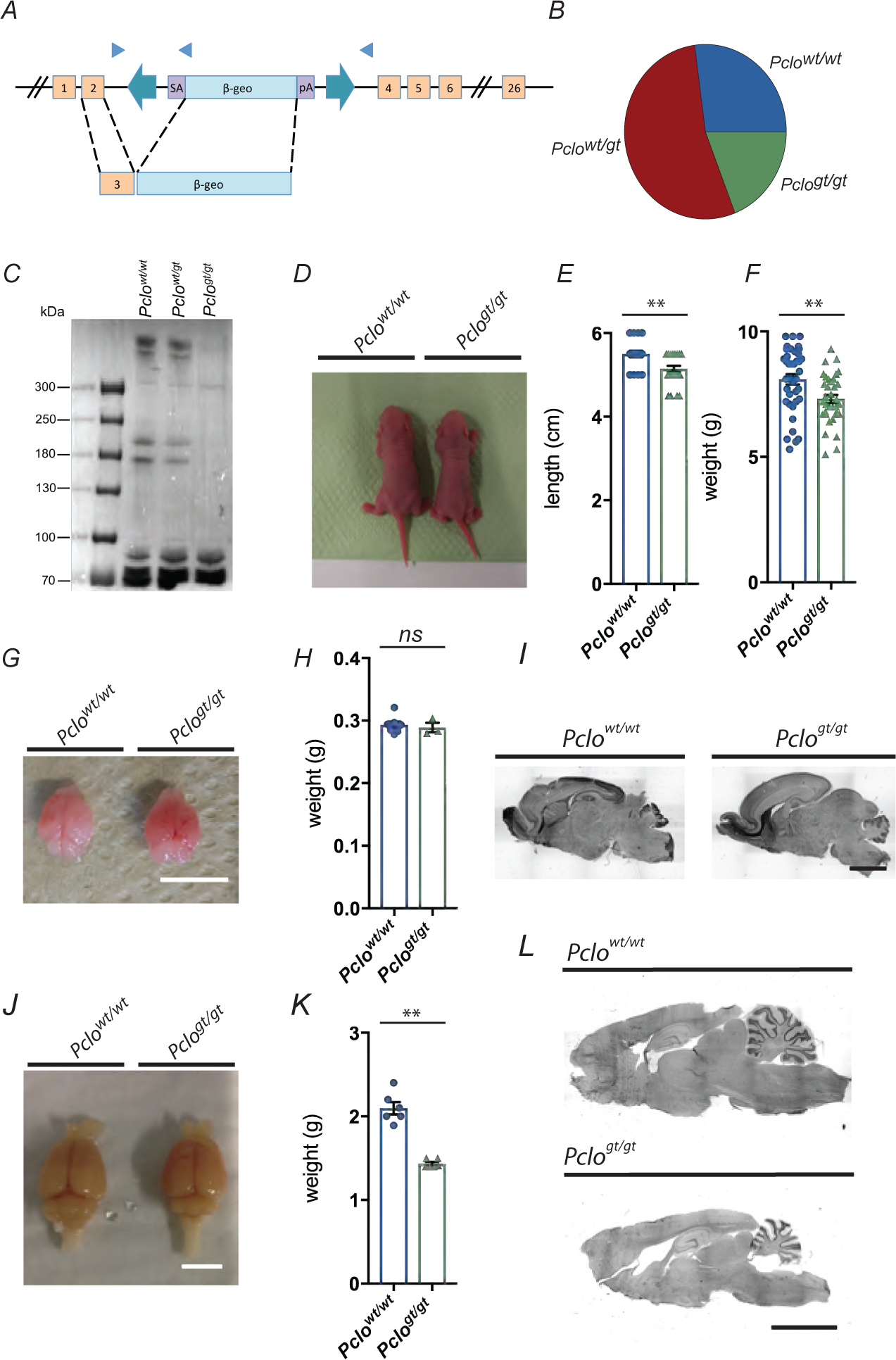
Generation of *Pclo^gt/gt^* mutant animals. A) Sleeping beauty transposon mutagenesis was used to generate gene trap (gt) piccolo knockout rats. The transposon element inserted into exon 3 of the piccolo genomic sequence and caused a stop in the reading frame. Adapted from Ackermann et al. (Ackermann et al., 2019). B) Pairs of heterozygous (*Pclo^wt/gt^*) males and females produced litters with Mendelian distribution. Pie chart demonstrates the birth rates of homozygous wildtype (*Pclo^wt/wt^*), homozygous gene trap mutation (*Pclo^gt/gt^*) and heterozygous (*Pclo^wt/gt^*) pups. C) Western blot analysis of brain lysates prepared from postnatal day 2 (P2) animals to confirm the loss of full-length Piccolo protein from the brain. A band of the Piccolo-corresponding size of 560 kDa is detectable in lysates prepared from *Pclo^wt/wt^* and *Pclo^wt/gt^* animals, but is absent in *Pclo^gt/gt^* brain lysates (data are representative of 3 independent experiments). However, smaller bands between 100 and 70 kDA are still present in *Pclo^gt/gt^* brain lysates. D-F) Image of postnatal day 1 (P1) littermates (D), E) Quantification of the body length of P0-P2 *Pclo^wt/wt^* and *Pclo^gt/gt^* pups (*Pclo^wt/wt^* = 5.5 cm ± 0.077, n = 23; *Pclo^gt/gt^* = 5.15 cm ± 0.070, n = 27; 6 independent litters; unpaired t-test, *p\*\** = 0.0014). F) Quantification of the body weight of P0-P2 *Pclo^wt/wt^* and *Pclo^gt/gt^* pups (*Pclo^wt/wt^* = 8.09 g ± 0.203, n = 39; *Pclo^gt/gt^* = 7.31 g ± 0.166, n = 35; 12 independent litters; unpaired t-test, *p\*\** = 0.0044). G-H) Image of brains dissected from P1 *Pclo^wt/wt^* and *Pclo^gt/gt^* pups (G). H) Quantification of the brain weight of P0-P2 *Pclo^wt/wt^* and *Pclo^gt/gt^* pups (*Pclo^wt/wt^* = 0.293 ± 0.00533, n = 7; *Pclo^gt/gt^* = 0.289 g ± 0.00758 n = 3; 3 independent litters; unpaired t-test, *p* = 0.683) I) Nissl stained sagittal sections from P2 rat brains show no overt differences between *Pclo^gt/gt^* and *Pclo^wt/wt^* pups. Scale bar = 0.5 cm. J-L) Image of 4% PFA-perfused brains from *Pclo^wt/wt^* and *Pclo^gt/gt^* animals at 3 months of age (I), K) Quantification of the brain weight showing that *Pclo^gt/gt^* brains are significantly lighter than *Pclo^wt/wt^* brains (*Pclo^wt/wt^* = 2.098 g ± 0.074, *Pclo^gt/gt^* = 1.435 g ± 0.021; n = 6, Mann-Whitney U test, *p\*\** = 0.0022). L) Nissl stained sagittal sections from 3 month-old rat brains reveal microcephaly in *Pclo^gt/gt^* compared to *Pclo^wt/wt^*. Note, ventricles are larger and cerebellum, pons, cerebrum and subcortical regions are smaller. Scale bars: 1 cm (G and K); 0.5 cm (I). Error bars represent SEM.

Intriguingly, the observed morphological changes in brains of 3 month-old *Pclo^gt/gt^* animals are remarkably similar to those reported for patients with PCH3, who exhibit microcephaly, a reduced size of the cerebrum, cerebellum and pons as well as larger ventricles (Maricich et al., 2011). As hypoplasia of the pons and cerebellum as well as a reduced thickness of the cortex are the most dramatic features of PCH3 patients, we examined in more detail changes occurring in these regions of *Pclo^gt/gt^* brains. Here, we found that the thickness of the cortex is significantly reduced in brains lacking Piccolo (Figure 2A and D). The area of the pons is dramatically reduced in *Pclo^gt/gt^* brain slices stained with Nissl (Figure 2B). As the density of neurons was not changed (Figure 2B, zoom), these data indicate a loss of the total number of neurons within the pons. This conclusion is supported by data from brainstem sections stained with antibodies against the synaptic vesicle (SV) protein VGluT1, prominently expressed in pons neurons, which reveals a smaller area occupied by these neurons (Figure 2C and E).

**Figure 2.**
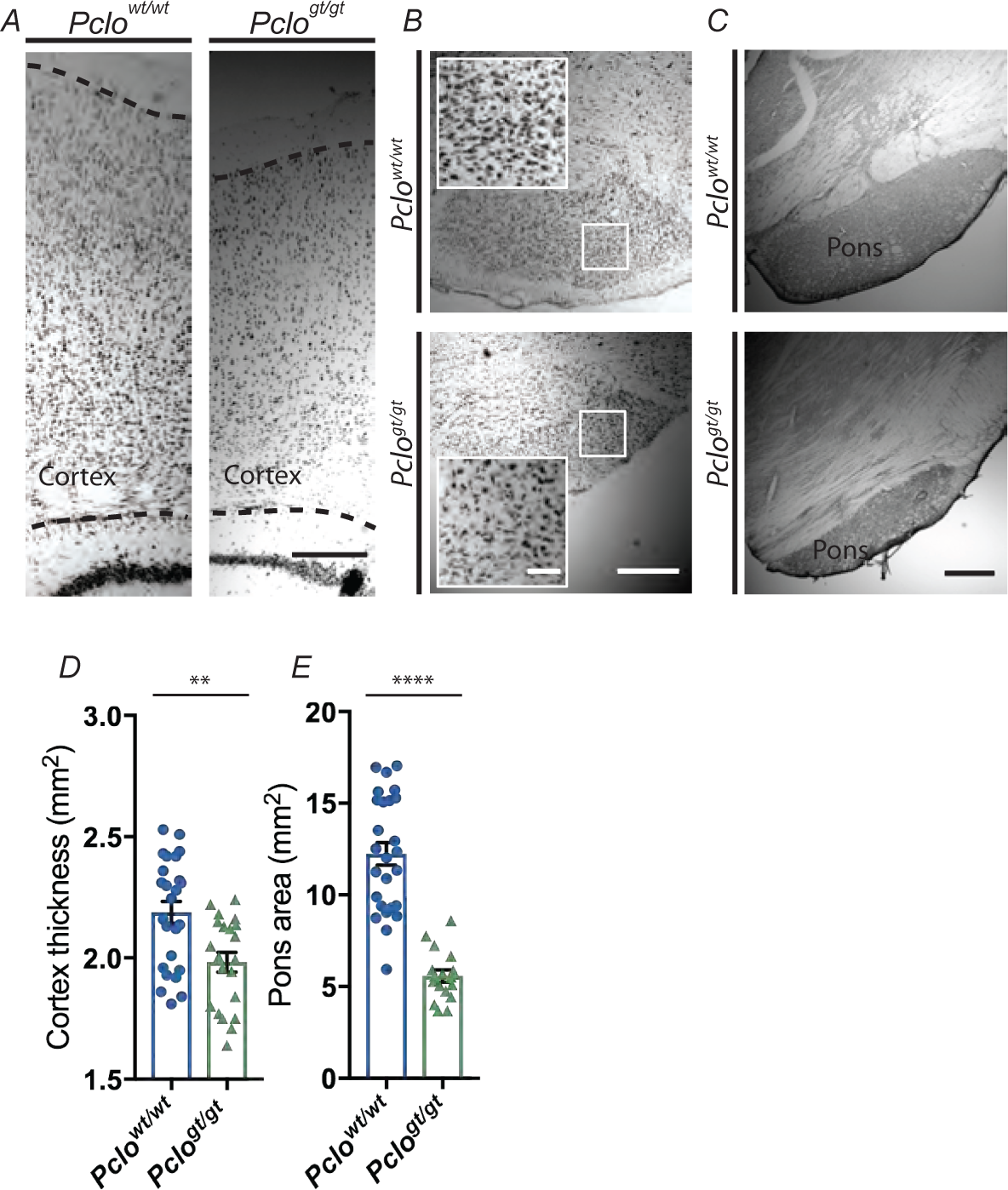
*Pclo^gt/gt^* rats show cortical thinning and a smaller pons area compared to *Pclo^wt/wt^* littermates. A-B) Nissl stained somatosensory cortex (indicated by dashed lines) (A) and the brainstem including pontine area (B) of *Pclo^wt/wt^* and *Pclo^gt/gt^* brains at 3 month of age. B) Zoom demonstrates that pontine neurons are of similar density in *Pclo^wt/wt^* and *Pclo^gt/gt^*. C) Pontine area visualized by staining with antibodies against VGluT1 and subsequent DAB conversion of *Pclo^wt/wt^* and *Pclo^gt/gt^* brains at 3 month of age. D) Quantification of the thickness of the somatosensory cortex (*Pclo^wt/wt^* = 2.18 mm ± 0.045, n = 25 brain sections; *Pclo^gt/gt^* = 1.98 mm ± 0.041, n = 21 brain sections; n = 3 independent experiments; unpaired t-test, *p\*\** = 0.0018). E) Quantification of the area of the pons (*Pclo^wt/wt^* = 12.24 mm^2^ ± 0.620, n = 26 brain sections; *Pclo^gt/gt^* = 5.58 mm^2^ ± 0.333, n = 17 brain sections; n = 3 independent experiments; unpaired t-test, *p**** <* 0.0001). Scale bars: 200 µm (A and C), 500 µm (B and C) and 100 µm (B, zoom). Error bars represent SEM.

Analysis of sagittal sections through the cerebellum of adult rats reveals that, whilst the cerebellum is smaller in *Pclo^gt/gt^* animals, the overall anatomy is not altered (Figure 3A). For example, there are no remarkable defects in foliation of *Pclo^gt/gt^* animals, with all lobes present and appearing to be formed normally at the vermis (Figure 3A and E). Our analysis of the granule cell layer (GCL), using DAPI to stain granule cell (GC) nuclei, reveals that this layer is significantly reduced in size in *Pclo^gt/gt^* compared to *Pclo^wt/wt^* controls (Figure 3A and C). However, this decrease was not associated with a proportional increase in GC packing density, as the number of GCs per GCL area is only very slightly increased (Figure 3B and D). These data indicate an overall loss of GCs in *Pclo^gt/gt^* cerebella. Conceptually, this is predicted to reduce the total number of GC parallel fibers innervating PC dendrites in the molecular layer (ML), a situation that could lead to a thinner ML and perhaps an altered packing density of Purkinje cells (PCs).

**Figure 3.**
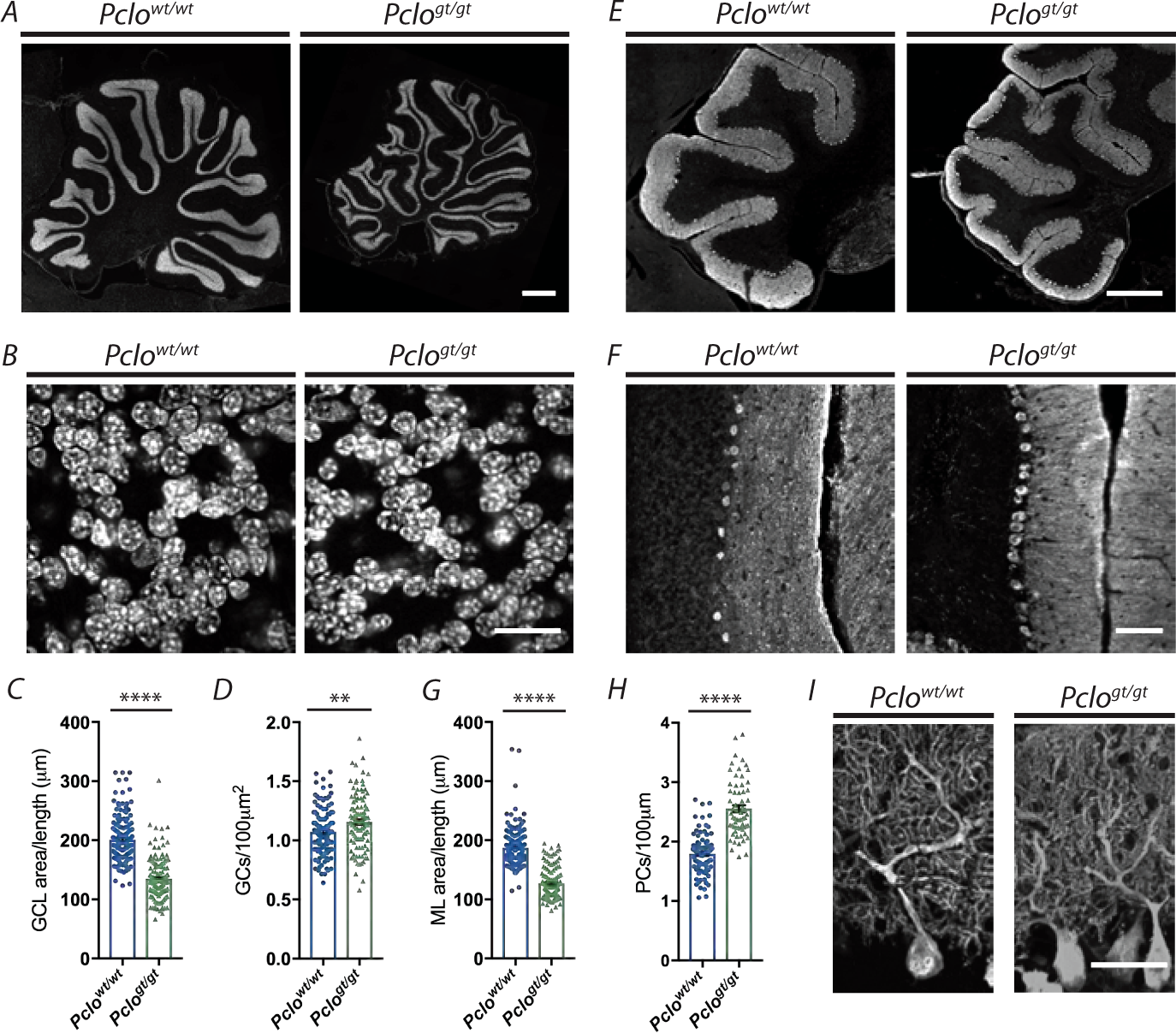
Morphometric comparison of *Pclo^gt/gt^* and *Pclo^wt/wt^* cerebella. A-D) Images of sagittal sections of *Pclo^wt/wt^* and *Pclo^gt/gt^* cerebella at 3 month of age. The densely packed granule cell layer (GCL) is visualized by DAPI staining. B) Higher magnification images of (A) demonstrating GC density in the GCL. C) Quantification of the GCL thickness (*Pclo^wt/wt^* = 200.8 µm ± 2.932, n = 160 lobes; *Pclo^gt/gt^* = 134.8 µm ± 2.859, n = 148 lobes; n = 4 independent experiments; Mann-Whitney U test, *p\*\*\*\** <0.0001). D) Quantification of the number of GCs per 100 µm^2^ (*Pclo^wt/wt^* = 1.702 ± 0.020, n = 111 images; *Pclo^gt/gt^* = 1.156 ± 0.022, n = 114 images; n = 3 independent experiments; unpaired t-test, *p\*\** = 0.0042). E-H) Images of sagittal sections of *Pclo^wt/wt^* and *Pclo^gt/gt^* cerebella at 3 months of age. Purkinje cells (PCs) stained with antibodies against Calbindin determine the molecular layer (ML) (lobes I-III shown). F) Higher magnification images of (E). Note the closer packing of PCs in *Pclo^gt/gt^* compared to *Pclo^wt/wt^*. G) Quantification of the ML thickness (*Pclo^wt/wt^* = 187.2 µm ± 2.719, n = 148 lobes; *Pclo^gt/gt^* = 127.2 µm ± 2.378, n = 125 lobes; n = 4 independent experiments; Mann-Whitney U test, *p\*\*\*\** < 0.0001). H) Quantification of the number of PCs per 100 µm length of PC layer (*Pclo^wt/wt^* = 1.797 ± 0.036; n = 89 lobes; *Pclo^gt/gt^* = 2.554 ± 0.058; n = 65 lobes; n = 3 independent experiments; unpaired t-test, *p\*\*\*\** < 0.0001). I) Images of sagittal sections stained with antibodies against Calbindin showing that PCs migrate correctly to their position in the ML and are correctly orientated. Scale bars: 1cm (A), 20 µm (B and J), 50 µm (I) and 200 µm (F). Error bars represent SEM. Data points represent images taken from lobes I, III, V, VII and IX; 4 sections per animal (B and D).

Sections immunostained with antibodies against Calbindin, which specifically labels PCs, reveals that the organization of the ML and the PC layer (PCL) appear to be intact and that the dendritic arbors of PCs are correctly orientated (Figure 3E, F and I). However, quantifying the average area per unit length of the ML reveals a dramatic reduction in the size of this layer in *Pclo^gt/gt^* brains (Figure 3G). Furthermore, cells in the PCL appear overcrowded in *Pclo^gt/gt^* cerebella (Figure 3E, F and I), a conclusion supported by data showing an increased packing density of PCs in *Pclo^gt/gt^* brains (Figure 3H).

Taken together, these data indicate a reduced number of GCs and a higher packing density of PCs in *Pclo^gt/gt^* cerebella compared to *Pclo^wt/wt^* controls.

### Loss of Piccolo alters climbing fiber and mossy fiber innervation in the cerebellum

Anatomically, the cerebellum receives its major excitatory afferents in the form of climbing fibers (CFs) and mossy fibers (MFs) that project from the inferior olive or the brainstem/spinal cord, respectively (Leto et al., 2016). Both form glutamatergic synapses but terminate in different layers of the cerebellum (Apps and Garwicz, 2005). For example, CFs form excitatory synapses onto the proximal branches of PC dendritic arbors, modulating the dynamic firing properties of PCs and motor learning (Hashimoto and Kano, 1998). In contrast, MFs terminate on the dendrites of GCs, which then provide direct excitatory input to PCs via their parallel fiber (PF) axons. Both also extend collaterals to the deep cerebellar nuclei before projecting into the cerebellar cortex (Shinoda et al., 1992).

In previous studies, we observed that Piccolo was present in the boutons of each of these excitatory synapses (Cases-Langhoff et al., 1996). This was confirmed by immunostaining cerebellar sections of *Pclo^wt/wt^* and *Pclo^gt/gt^* with antibodies against Piccolo and the SV protein VGlut1 (Fig. 4A and B). This revealed that Piccolo immuno-reactivity was indeed present at each of these synaptic types and that this immuno-reactivity was lost in cerebella from *Pclo^gt/gt^* animals. To explore whether deficiencies in either could contribute to the anatomical and functional changes in the cerebellum, we initially analyzed potential differences in the CF input into the ML. Synaptic input from CFs onto PC dendrites, immuno-positive for Calbindin, was visualized with antibodies against VGluT2 (Miyazaki et al., 2003). Here our analysis of sagittal sections revealed the presence of a large number of VGluT2 positive puncta decorating Calbindin positive dendrites in *Pclo^gt/gt^* and *Pclo^wt/wt^* sections (Figure 4C and D). These data indicate that the loss of Piccolo does not affect the ability of CFs to project into the ML and form synapses with PC soma and dendrites. Qualitatively, VGluT2 positive puncta in *Pclo^wt/wt^* and *Pclo^gt/gt^* sections were of similar size and beautifully decorated both primary and tertiary PC dendrites, though the total number of puncta appeared more numerous in *Pclo^gt/gt^* sections. Quantification of the total area of VGluT2 per ML supports this impression (Figure 4D). Additionally, we compared the distribution of the CFs to the synaptic inputs from GCs onto PCs, the parallel fiber (PF) axons which project into the ML. The latter synapses were identified with antibodies against VGluT1 (Miyazaki et al., 2003). Although the ML is thinner in *Pclo^gt/gt^* cerebella (Figure 3A), we observed no overt changes in the intensity or distribution of VGluT1 positive puncta throughout the ML of *Pclo^gt/gt^* animals compared to *Pclo^wt/wt^* controls (Figure 4C). This implies that PF axons project normally and form a robust number of synapses with PCs. Given that fewer GCs are formed in *Pclo^gt/gt^* cerebella (Figure 3A and B), we postulate that the thinner ML is most likely due to fewer PFs and less total synaptic input on PC dendrites.

**Figure 4.**
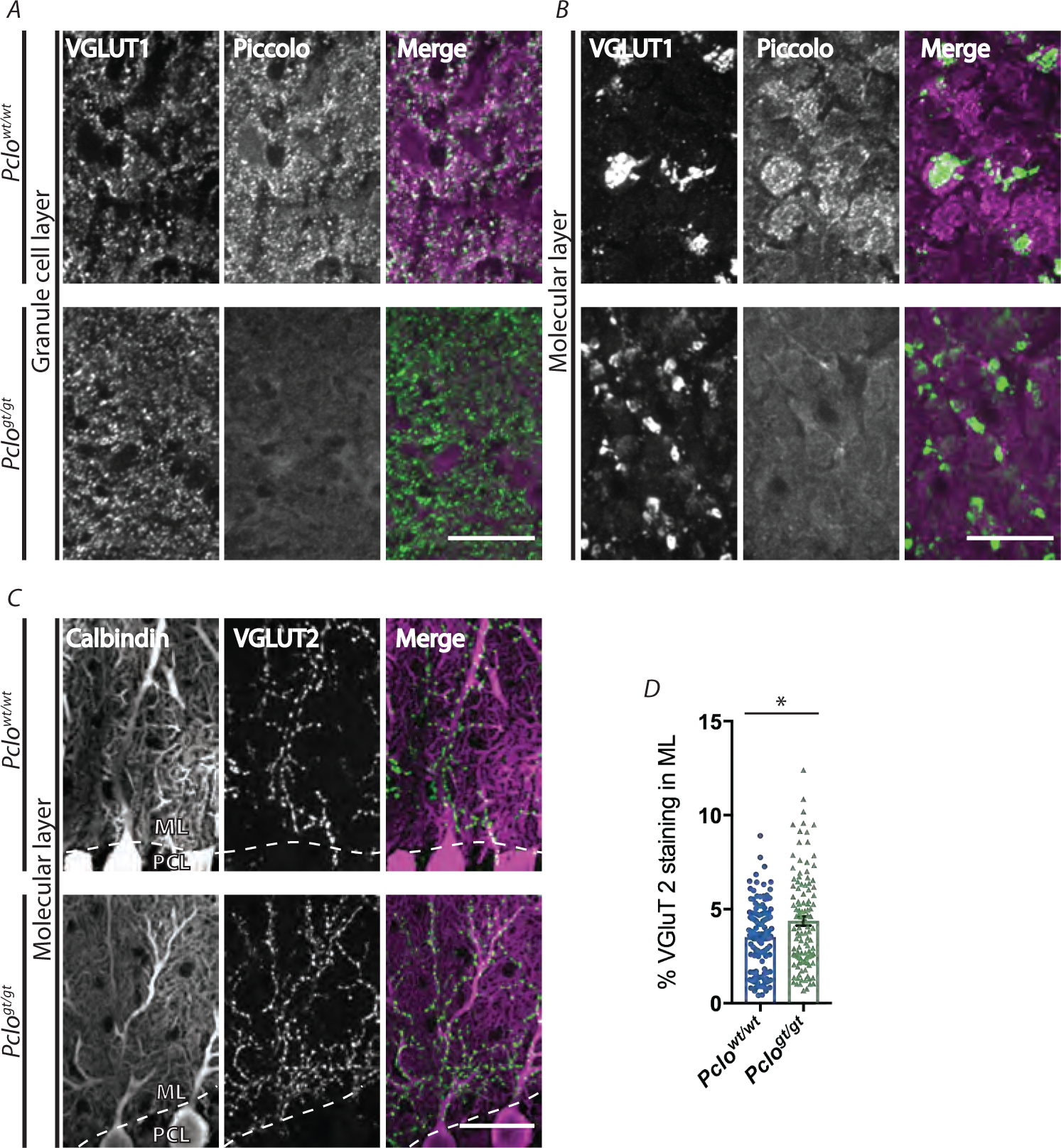
Abberant climbing fiber innervation of Purkinje cells in *Pclo^gt/gt^* rats compared to *Pclo^wt/wt^* littermates. A and B) Images of sagittal sections of *Pclo^wt/wt^* and *Pclo^gt/gt^* cerebella at 3 months of age stained with antibodies against VGluT1 and Piccolo demonstrate the loss of Piccolo in the ML (A) and the GCL (B) in *Pclo^gt/gt^* rats. C) Images of sagittal sections of *Pclo^wt/wt^* and *Pclo^gt/gt^* cerebella at 3 months of age stained with antibodies against Calbindin and VGluT2. Note that the climbing fiber synapses, immuno-positive for VGluT2, are increased in the ML of *Pclo^gt/gt^* cerebella compared to *Pclo^wt/wt^* controls (C and D). When Piccolo is absent, parallel fiber synapses in the ML - immuno-positive for VGluT1 – do not appear overtly different between *Pclo^wt/wt^* and *Pclo^gt/gt^* (A). However, MF and CF synapses are altered (4B-D, Figure 5). D) Quantification of the percentage of the ML (indicated by dashed lines) immuno-positive for VGluT2 from (C) (*Pclo^wt/wt^* = 3.521 ± 0.160, n = 128 images; *Pclo^gt/gt^* = 4.377 ± 0.241, n = 112 images; n = 3 independent experiments; Mann-Whitney U test, *p\** = 0.0278). Scale bars: 20 µm. Error bars represent SEM. Data points represent images taken from lobes I, III, V, VII and IX; 4 sections per animal.

GCs are known to receive their excitatory input from MFs arising from afferent axons from a number of distinct nuclei in the brainstem including the pontine nuclei (Sillitoe, 2012). These collaterals form large glomerular structures with multiple AZs, forming a rosette of synapses with claws from dendrites of multiple GCs (Jakab and Hamori, 1988; Xu-Friedman and Regehr, 2003). Given the smaller size of the pons, we thus explored whether the boutons from the remaining cells properly reached the cerebellum and formed robust MF terminals. As afferent fibers from the pons primarily innervate cerebellar lobes VI to IX, we examined sagittal sections of these lobes stained with antibodies against the somatodendritic marker MAP2 and VGluT2 in lobe VII. In *Pclo^wt/wt^* sections, multiple large VGluT2 positive puncta are seen packed tightly together within a dense meshwork of MAP2 positive dendrites projecting from a ring of GCs. These puncta represent subclusters of SVs with large 100-200 µm^3^ terminals (Jakab and Hamori, 1988). In sections from *Pclo^gt/gt^* animals, each bouton appeared smaller in size and less organized, though they still appear to contact GC dendrites (Figure 5A). This impression is further supported by sections immunostained with antibodies against VGluT1 and VGluT2. In these images, distinct VGluT1 and/or VGluT2 positive clusters are observed in both *Pclo^wt/wt^* and *Pclo^gt/gt^* animals, consistent with individual boutons or glomeruli (Figure 5B). Here again, MF terminals from *Pclo^gt/gt^* animals appeared to have much smaller VGluT1 or VGluT2 positive clusters (Figure 5B, zoom). Quantifying the average size of the VGluT1 and VGluT2 positive clusters revealed that clusters of *Pclo^wt/wt^* cerebella were more than double the size compared to the same lobes in *Pclo^gt/gt^* animals (Figure 5B and C). However, this effect was not lobe-specific as it was observed in all lobes and not just in those receiving pontocerebellar afferents (Figure 5B). These findings suggest that defects in the formation of large robust MF glomeruli in *Pclo^gt/gt^* cerebella is a common feature shared by MF afferents arising from the pons and other brainstem nuclei, and may reflect aberrant signaling during development between GCs and these neurons.

**Figure 5.**
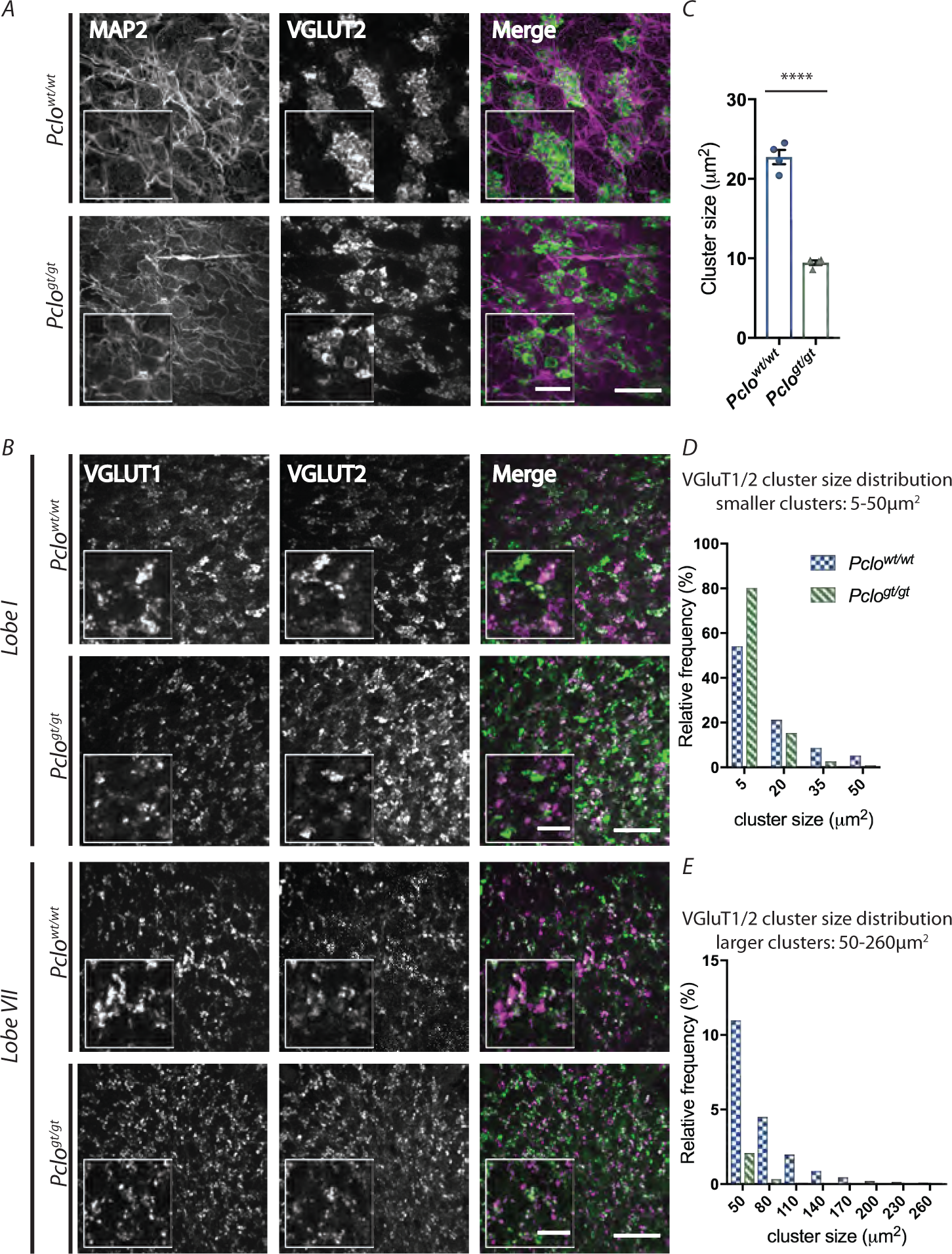
Cerebella from *Pclo^gt/gt^* animals have smaller mossy fiber rosettes. A and B) Images of sagittal sections of *Pclo^wt/wt^* and *Pclo^gt/gt^* cerebella at 3 month of age stained with antibodies against VGluT2, which is highly expressed at mossy fiber (MF) boutons, and the somatodendritic marker MAP2 (A) or VGluT2 and VGluT1 (B). A) GCs extend their dendrites into MF boutons in the *Pclo^wt/wt^* condition. However, in *Pclo^gt/gt^*, whilst GC dendrites are still juxtaposed to VGluT2-positive boutons, the boutons are smaller and therefore the arrangement is less organized. B) Presynaptic MF glomeruli from lobes I (upper panel) and VII (lower panel) are visualized by VGluT1 and VGluT2. Note that the reduction in MF size is consistent regardless of the lobe. Note also that rosettes are generally labeled with either VGluT1 or VGluT2 and occasionally with both markers consistent with them being innervated by a single synaptic input from different neuronal cell types. C) Quantification of the size of VGluT1/VGluT2 clusters (*Pclo^wt/wt^* = 22.73 µm^2^ ± 0.896, n = 4 animals; *Pclo^gt/gt^* = 9.46 µm^2^ ± 0.2899, n = 4 animals; unpaired t-test, *p\*\*\*\** < 0.0001). D-E) Histograms to show the distribution of puncta sized 5-50 µm^2^ (D) and 50-260 µm^2^ (E). The shift of the data indicates *Pclo^gt/gt^* MFs have more smaller puncta (5 µm^2^), whereas *Pclo^wt/wt^* MFs have more larger puncta (up to 260 µm^2^). Scale bars: 50 µm (A and B) and 20 µm (B, zoom). Error bars represent SEM. Data points represent average puncta size per animal from images taken from lobes I, III, V, VII and IX, 4 sections per animal (C, D and E).

Quantitatively, the spread of distribution of VGluT1 and VGluT2 cluster sizes is far more shifted towards smaller cluster sizes in *Pclo^gt/gt^*, with mutant cerebellar displaying approximately 45 % more small synaptic clusters of 5 µm^2^ in size than *Pclo^wt/wt^* counterparts (Figure 5D). At the larger end of the scale, cluster sizes over 50 µm^2^ were much more frequent for *Pclo^wt/wt^* than *Pclo^gt/gt^* cerebella (Figure 5E).

In addition to excitatory input, MF glomeruli are also modulated by GABAergic inhibition via cerebellar Golgi cells, which offer regulatory feedback to the complex, as they themselves are excited by GCs (Maex and De Schutter, 1998). In principle, smaller MFs terminals seen in Piccolo KO animals could represent less excitatory input into GCs. This may also reduce excitatory drive onto PCs via PFs from GCs, as well as perhaps inhibitory drive via the Golgi cells. One of the dominant receptors mediating inhibitory input from Golgi cells to GCs are the α6 subunit-expressing GABA_A_ receptors, which are highly concentrated within MF glomeruli (Nusser et al., 1996). Sagittal sections immunostained with antibodies against VGluT2 and α6 subunits revealed that the GABA_A_ α6 receptor subunit is only weakly expressed in *Pclo^gt/gt^* MF rosettes, whereas in *Pclo^wt/wt^*, it nicely localizes to the synaptic complex (Figure 6A). Antibody staining of the granule cell layer demonstrates higher intensity of GABA_A_ α6 antibody staining in *Pclo^wt/wt^* compared to *Pclo^gt/gt^* (Figure 6B). However, knockout models of GABA_A_α6 (Homanics et al., 1997) do not display PCH3 phenotypes or alterations in the anatomy of the cerebellum. Therefore, GABA_A_α6 downregulation can be attributed to Piccolo loss and not for the PCH3 phenotypes.

**Figure 6.**
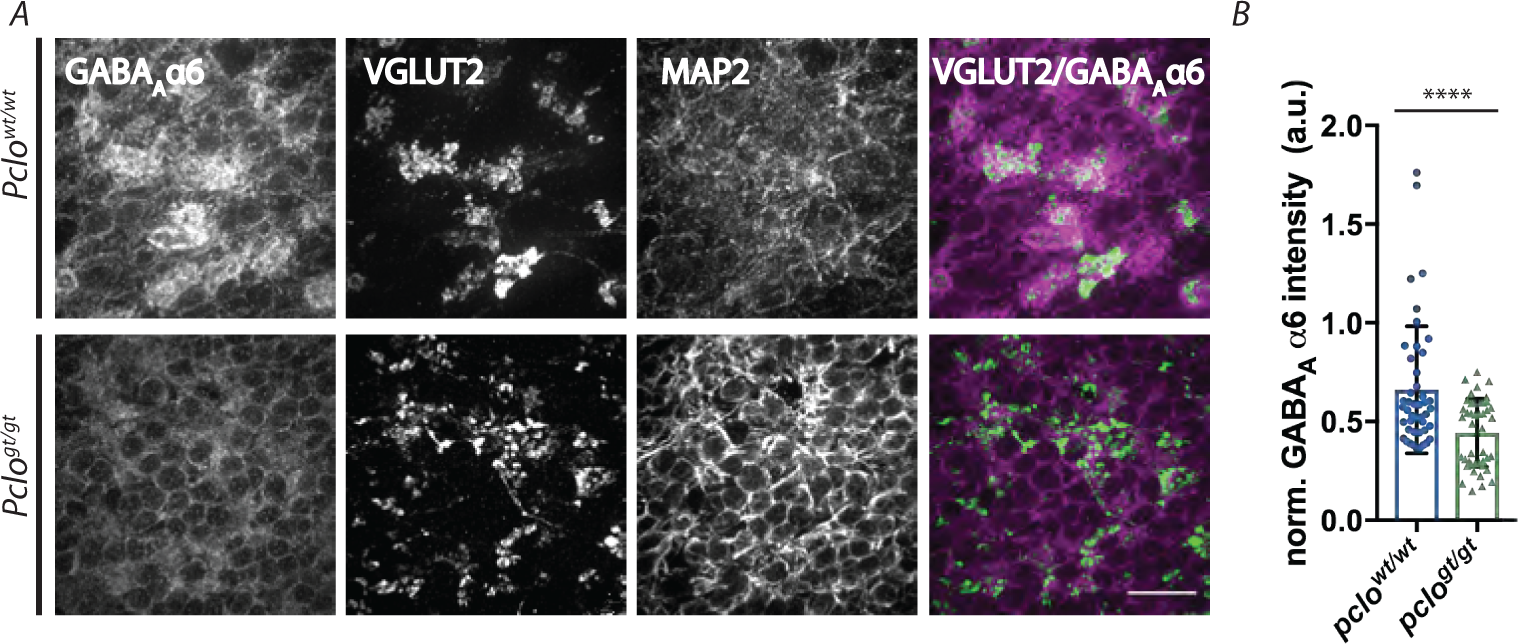
GABA_A_α6 subunit expression is lower in *Pclo^gt/gt^* than in *Pclo^wt/wt^*. A) Representative images of sagittal sections of *Pclo^wt/wt^* and *Pclo^gt/gt^* cerebella at 3 months of age stained with antibodies against the GABA_A_ subunit α6,VGluT2 and MAP2. Note the decreased levels of GABA_A_α6 in the GC layer of *Pclo^gt/gt^* animals compared to *Pclo^wt/wt^* controls, quantified in B). B) Quantification of GABA_A_ subunit α6, measured by the average intensity of antibody staining in images taken from the GC of the cerebellum, normalized to MAP2 antibody intensities for the same image (*Pclo^wt/wt^* = 0.661 ± 0.0479 arbitrary units (a.u.), n = 45 images from 3 individual animals; *Pclo^gt/gt^* = 0.443 ± 0.26 a.u., n = 44 images from 3 individual animals; Mann-Whitney U test, *p*\*\*\* = 0.0009). Scale bars: 20 µm. Error bars represent SEM. Data points represent images taken from 4 sections per animal.

These data suggest that the loss of Piccolo not only affects the size of MF inputs into the cerebellum, but also gene expression and therefore inhibitory drive within each glomerulus, a condition that could relate to the maturation of these structures and/or their functionality, a situation that could adversely affect cerebellar function.

### Ultrastructural analysis of mossy fiber glomeruli in *Pclo^gt/gt^* cerebellum

The reduced area of VGluT1 and VGluT2 clusters within *Pclo^gt/gt^* MF terminals (Figure 6 could be due to a reduction in size of MF glomeruli themselves and/or in the number of VGluT1/2 positive SVs per bouton. To explore these options, we investigated MF glomeruli in *Pclo^wt/wt^* and *Pclo^gt/gt^* cerebella using electron microscopy (EM). Analysis of ultrathin cerebellar brain sections from 3 month-old rats revealed that the average size of *Pclo^gt/gt^* glomeruli was significantly smaller than *Pclo^wt/wt^* glomeruli (Figure 7A and C). Furthermore, the complexity of the glomeruli indicated by a P2A value, measuring the ratio of perimeter per area, was significantly reduced (Figure 7E). The size of the presynaptic area was also reduced in *Pclo^gt/gt^* cerebellar sections (Figure 7B and D). However, the number of active zones (AZs) present at each glomerulus was still proportional to their size, as *Pclo^wt/wt^* and *Pclo^gt/gt^* had a similar number of AZs per glomerular area (Figure 7B and F). Given that *Pclo^gt/gt^* boutons are smaller, this indicates that the overall output of the MF glomeruli could be reduced in *Pclo^gt/gt^* cerebellum. Intriguingly, we also noticed an accumulation of clathrin-coated vesicles (CCVs) in the terminals of *Pclo^gt/gt^* MFs (Figure 7G). This phenotype resembles recent findings from hippocampal synapses from *Pclo^gt/gt^* animals, which revealed defects in the formation of endosomal membranes and an overall reduction in SV number (Ackermann et al., 2019) and suggest possible changes in the recycling of SVs within MF boutons post fusion with the plasma membrane.

**Figure 7.**
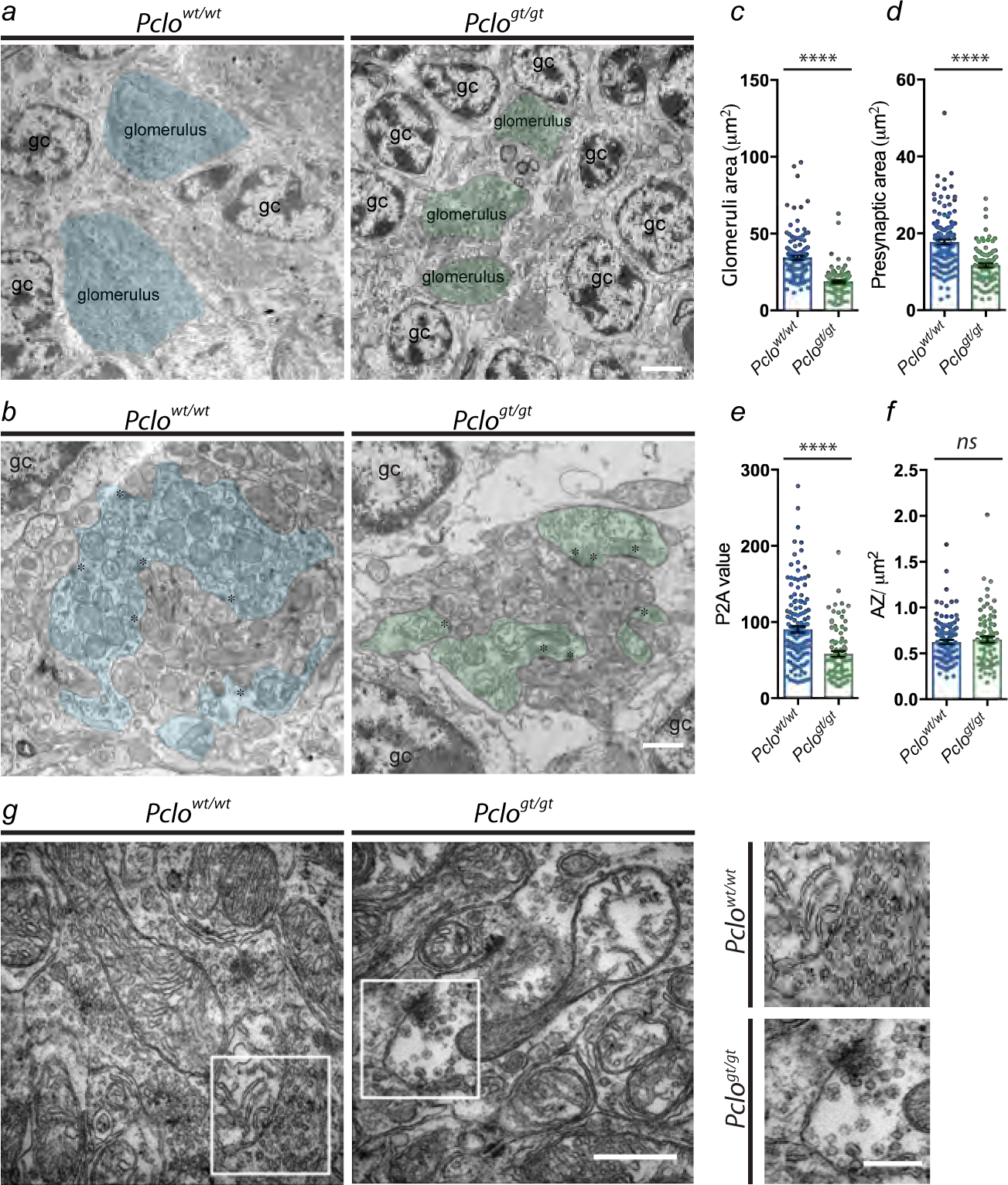
Glomerular rosettes are smaller and less complex in *Pclo^gt/gt^*. A, B and G) Electron microscopy images of the GCL of *Pclo^wt/wt^* and *Pclo^gt/gt^* cerebella at 3 months of age. Granule cells are indicated by ‘gc’ (A and B), cerebellar glomeruli are highlighted in blue (*Pclo^wt/wt^*) and green (*Pclo^gt/gt^*) (A) and the presynaptic terminals of cerebellar MF are highlighted in blue (*Pclo^wt/wt^*) and green (*Pclo^gt/gt^*) (B). Asterisks mark active zones (AZs) (B) and the zoom in (G) emphasizes the presence of more clathrin-coated vesicles (CCVs) in *Pclo^gt/gt^* MF boutons compared to than *Pclo^wt/wt^*. (C-F) Quantification of the size of glomeruli (C), the size of the MF presynapse (D), the complexity (squared perimeter/presynaptic area) of the MF bouton (E) and the density of AZs (F). Note the strong decrease in glomerulus and MF bouton size (C and D) (C) (*Pclo^wt/wt^* = 34.59 µm^2^ ± 1.287, n = 130 images; *Pclo^gt/gt^* = 18.82 µm^2^ ± 0.853, n = 103 images; n = 3 independent experiments; Mann-Whitney U test, *p\*\*\*\** < 0.0001. (D) (*Pclo^wt/wt^* = 17.73 µm^2^ ± 0.603, n = 141 images; *Pclo^gt/gt^* = 11.69 µm^2^ ± 0.497, n = 95 images; n = 3 independent experiments; Mann-Whitney U test, *p\*\*\*\** < 0.0001). (E) (*Pclo^wt/wt^* = 90.5 ± 4.089, n = 141 images; *Pclo^gt/gt^* = 58.1 ± 3.818, n = 84 images; n = 3 independent experiments; Mann-Whitney U test, *p\*\*\*\** < 0.0001). (F) (*Pclo^wt/wt^* = 0.625 ± 0.021, n = 121 images; *Pclo^gt/gt^* = 0.652 ± 0.032, n = 84 images; n = 3 independent experiments; Mann-Whitney U test, *p* = 0.7390). Scale bars: 2.5 µm (A), 1µm (B), 500 nm (G) and 250 nm (G, zoom). Error bars represent SEM.

### Piccolo loss alters GC properties and mossy fiber to GC synaptic transmission

The anatomical and morphological changes observed in MF boutons lacking Piccolo are predicted to not only represent altered afferent input into the cerebellum from the pons and other brainstem nuclei, but also altered cerebellar function. As an initial test of this hypothesis, we performed whole-cell current clamp recordings of cerebellar GCs from acute P90 rat cerebellar slices. A two-photon image of a typical cerebellar GC from *Pclo^gt/gt^* filled with ATTO dye reveals a normal radial arrangement of its dendrites as they project their claws into MF glomeruli (Figure 8A). An analysis of the intrinsic biophysical properties of these cells revealed that the GC properties differed between *Pclo^gt/gt^* and *Pclo^wt/wt^* animals. Specifically, no changes were detected in either the capacitance or the membrane potential of these cells, but the input resistance was significantly increased (Figure 8B). Since these experiments were performed in the presence of GABA_A_ receptor blockers, the decreased shunting inhibition mediated by tonic activation of α6-subunit-containing GABA_A_ receptors (Brickley et al., 1996; Nusser et al., 1998) is expected to further increase the difference in input resistance. Yet, there was also no change in the amplitude, threshold of activation or duration of action potentials fired by these cells (Figure 8B), indicating unaltered active membrane properties. Examining the frequency and amplitude of spontaneous miniature excitatory postsynaptic currents (mEPSCs) of these GCs revealed a dramatic increase in the frequency of these events in *Pclo^gt/gt^* slices with no change in mEPSC amplitudes (Figure 8C). These data suggest that on average each excitatory synapse formed on to these GC dendrites has normal levels of postsynaptic AMPA-type glutamate receptors. The change in frequency could either be due to an increase in the number of MF boutons contacting GC dendrites and/or an increase in the release probability of MF boutons. Consistently, the average amplitude of the evoked excitatory postsynaptic currents (EPSCs) was increased in *Pclo^gt/gt^* animals, with no change in the weighted time constant (τ_w_) (Figure 8D). Given the amplitudes of the mEPSCs are not changed, these results suggest that there is a higher number of synaptic connections between MF boutons and the dendrites of GCs, consistent with the larger number of smaller SV clusters/rosette in *Pclo^gt/gt^* animals (Figure 6). Taken together, these data indicate that, in addition to changes in cell number, the loss of Piccolo in the cerebellum and brainstem is associated with changes in the morphology and function of GCs and their mossy fiber input.

**Figure 8.**
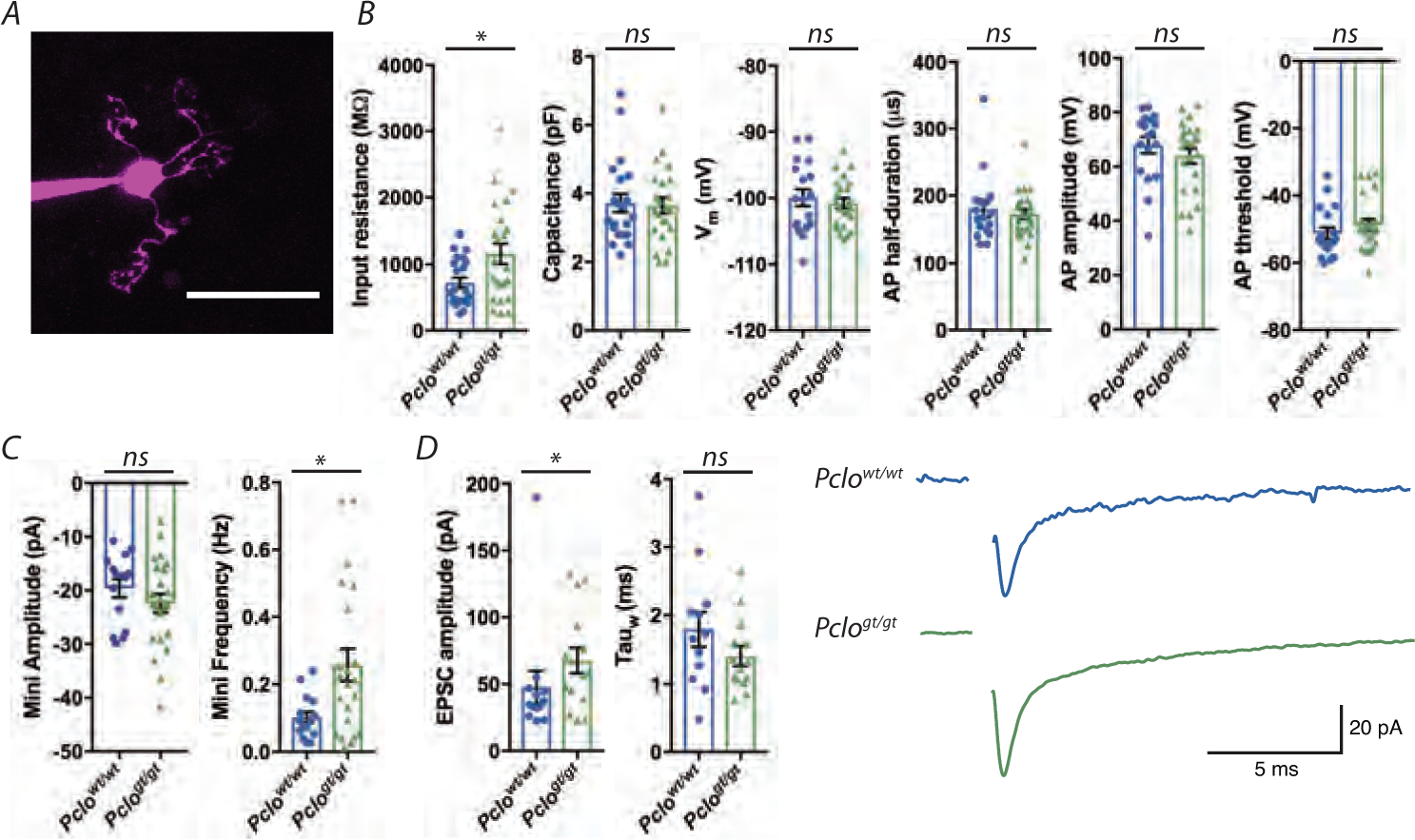
Physiological assessment of mossy fiber boutons. A) Example Two-photon image of a cerebellar granule cell (GC) from a *Pclo^gt/gt^* rat filled with ATTO dye. B) Average data of biophysical properties of GCs for *Pclo^wt/wt^* and *Pclo^gt/gt^* rats. The input resistance of GCs was higher in *Pclo^gt/gt^* compared to *Pclo^wt/wt^* (Pclowt/wt = 722.6 MΩ ± 76.24, n = 22 cells; *Pclo^gt/gt^* = 1160 MΩ ± 154.9, n = 23 cells; n = 3 rats per genotype; Mann-Whitney U test, *p*\* = 0.0462). Whereas no differences were found in capacitance (*Pclo^wt/wt^* = 3.717 pF ± 0.269, n = 21 cells; *Pclo^gt/gt^* = 3.643 pF ± 0.231, n = 23 cells; n = 3 rats per genotype; Mann-Whitney U test, *p* = 0.912), resting membrane potential (Vm) (*Pclo^wt/wt^* = −99.96 mV ± 1.261, n = 18 cells, *Pclo^gt/gt^* = −100.8 mV ± 0.786, n = 23 cells; n = 3 rats per genotype; Mann-Whitney U test, *p* = 0.612), the half-duration of the action potential (*Pclo^wt/wt^* = 179.7 µs ± 11.84, n = 18 cells; *Pclo^gt/gt^* = 172.2 µs ± 7.366, n = 23 cells; n = 3 rats per genotype; Mann-Whitney U test, *p* = 0.866), the amplitude of the action potential (*Pclo^wt/wt^* = 67.85 mV ± 3.016, n = 18 cells; *Pclo^gt/gt^* = 63.92 mV ± 2.761, n = 23 cells; n = 3 rats per genotype; Mann-Whitney U test, *p* = 0.291) and the voltage threshold to elicit an action potential (*Pclo^wt/wt^* = −51.27 mV ± 1.748, n = 18 cells; *Pclo^gt/gt^* = −48.6 mV ± 1.659, n = 23 cells; n = 3 rats per genotype; Mann-Whitney U test, *p* = 0.162). C) Miniature excitatory postsynaptic currents from *Pclo^gt/gt^* GCs were not different in their amplitude (*Pclo^wt/wt^* = −19.62 pA ± 1.682, n = 15 cells; *Pclo^gt/gt^* = −22.44 pA ± 1.765, n = 23 cells; n = 3 rats per genotype; Mann-Whitney U test, *p* = 0.286) but in their frequency (*Pclo^wt/wt^* = 0.102 Hz ± 0.0167, n = 15 cells; *Pclo^gt/gt^* = 0.257 Hz ± 0.0481, n = 22 cells; n = 3 rats per genotype; Mann-Whitney U test, *p*\* = 0.0392). D) Excitatory postsynaptic currents from GCs measured after stimulation of single mossy fibers were increased in *Pclo^gt/gt^* compared to *Pclo^wt/wt^* (*Pclo^wt/wt^* = 47.58 pA ± 12.12, n = 13 cells; *Pclo^gt/gt^* = 67.62 pA ± 9.64, n = 15 cells; n = 3 rats per genotype; Mann-Whitney U test, *p*\* = 0.0356), whereas the decay of the EPSC was not alerted (*Pclo^wt/wt^* = 1.79 ms ± 0.258, n = 12 cells; *Pclo^gt/gt^* = 1.404 ms ± 0.141, n = 14 cells; n = 3 rats per genotype; Mann-Whitney test U, *p*\* = 0.231). Right hand panel: example traces of evoked EPSCs, as quantified in D), in response 1 Hz stimulation in the presence of 20 µM SR95531 and 40 µM D-(2R)-amino-5-phosphonovaleric acid (D-APV). Scale bar = 20 µm (A). Error bars represent SEM. Data points represent individual cells from 3 rats per genotype.

### Behavioral and motor defects in Piccolo knockout rats

The motor difficulties reported (Zelnik et al., 1996; Durmaz et al., 2009) in humans with PCH3 and the anatomical changes observed in the cerebellum and brainstem of rats lacking Piccolo predict altered motor function in these animals. To test this hypothesis, *Pclo^wt/wt^*, *Pclo^wt/gt^* and *Pclo^gt/gt^* rats were monitored for their motor abilities. In rotarod tasks, *Pclo^gt/gt^* rat performance was significantly reduced compared to *Pclo^wt/wt^* and *Pclo^wt/gt^* rats (Figure 9A). Specifically, while *Pclo^wt/wt^* and *Pclo^wt/gt^* rats exhibited increasing performance levels regarding the ability to stay on the rotarod over time, *Pclo^gt/gt^* rats showed no indication of being able to adapt to the task. *Pclo^gt/gt^* rats were less adept at staying on the task apparatus once rod rotation was initiated (Figure 9A). Intriguingly, no differences in forelimb grip strength were scored between Piccolo genotypes (Figure 9B), indicating that *Pclo^gt/gt^* rats showed lack of motivation and/or impaired coordination.

**Figure 9.**
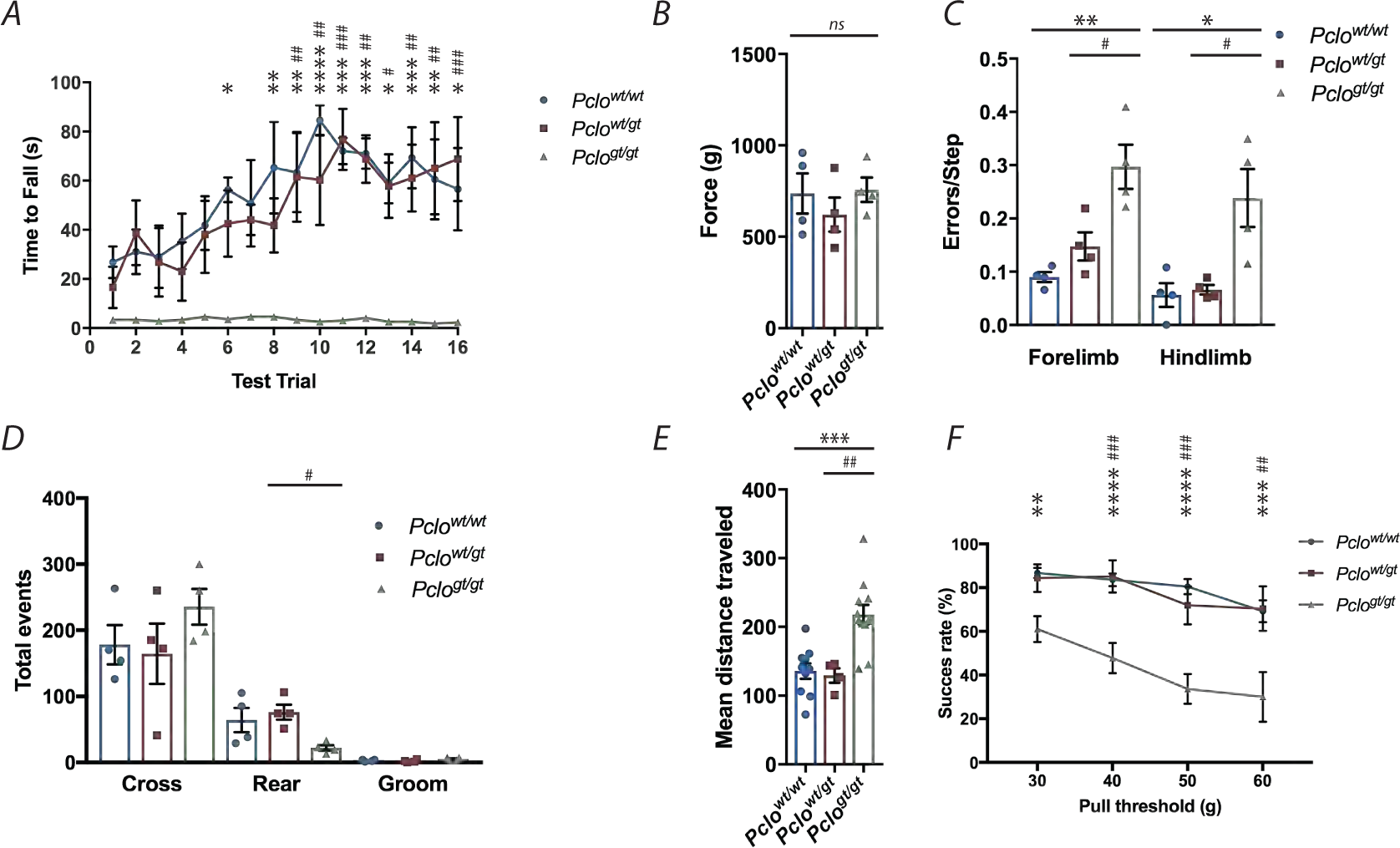
Behavioral outcome of Piccolo loss of function resembles PCH3 symptoms. A) Rotarod performance for *Pclo^wt/wt^*, *Pclo^wt/gt^* and *Pclo^gt/gt^* rats for 16 trials over 4 days. *Pclo^gt/gt^* rats fell significantly faster than *Pclo^wt/wt^* on trials 6 and 8 and faster than both *Pclo^wt/wt^* and *Pclo^wt/gt^* on trials 9-16 (trial 6: *Pclo^wt/wt^* = 56.25 s ± 5.023, *Pclo^gt/gt^* = 3.5 ± 0.5, *p\** = 0.0293; trial 8: *Pclo^wt/wt^* = 65.25 s ± 18.688, *Pclo^gt/gt^* = 4.5 ± 0.957, *p\*\** = 0.0042; trial 9: *Pclo^wt/wt^* = 63.25 s ± 16.163, *Pclo^wt/gt^* = 61.5 s ± 18.319, *Pclo^gt/gt^* = 3.25 ± 0.75, *p\*\** = 0.0051, *p*^##^ = 0.0079 (where * **=** *p* value between *Pclo^wt/w^* and *Pclo^gt/gt^* and ^#^ = *p* value between *Pclo^wt/wt^* and *Pclo^wt/gt^*); trial 10: *Pclo^wt/wt^* = 84.5 s ± 6.225, *Pclo^wt/gt^* = 60.25 s ± 18.396, *Pclo^gt/gt^* = 2.5 ± 0.5, *p\*\*\*\** = <0.0001, *p^##^* = 0.0089; trial 11: *Pclo^wt/wt^* = 72 s ± 5.339, *Pclo^wt/gt^* = 76.75 s ± 12.419, *Pclo^gt/gt^* = 3 ± 0.408, *p\*\*\** = 0.0005, *p^###^* =0.001; trial 12: *Pclo^wt/wt^* = 71 s ± 6.178, *Pclo^wt/gt^* = 68.75 s ± 9.681, *Pclo^gt/gt^* = 4 ± 0.707, *p\*\*\** = 0.0008, *p^##^* = 0.0015; trial 13: *Pclo^wt/wt^* = 59 s ± 8.297, *Pclo^wt/gt^* = 57.75 s ± 12.99, *Pclo^gt/gt^* = 2.5 ± 0.5, *p\** = 0.0121, *p^#^* = 0.0164; trial 14: *Pclo^wt/wt^* = 69.25 s ± 12.479, *Pclo^wt/gt^* = 61 s ± 13.638, *Pclo^gt/gt^* = 2.5 ± 0.645, *p\*\*\** = 0.0009, *p^##^* = 0.0074; trial 15: *Pclo^wt/wt^* = 60.5 s ± 16.297, *Pclo^wt/gt^* = 65 s ± 18.757, *Pclo^gt/gt^* = 1.75 ± 0.479, *p\*\** = 0.0070, *p^##^* = 0.0022; trial 16: *Pclo^wt/wt^* = 56.5 s ± 16.775, *Pclo^wt/gt^* = 68.75 s ± 17.109, *Pclo^gt/gt^* = 2.25 ± 0.25, *p\** = 0.0207, *p^###^* = 0.009; n = 3 animals per genotype; two-way ANOVA with Bonferroni correction). B) Grip strength task for *Pclo^wt/wt^*, *Pclo^wt/gt^* and *Pclo^gt/gt^* rats for 9 trials over 2 days. No differences were found for forelimb grip strength between the groups. (*Pclo^wt/wt^* = 737.2 ± 109.9; *Pclo^wt/gt^* = 621.2 ± 93.46, *Pclo^gt/gt^* = 757.5 ± 66.74; n = 4 rats per genotype, one-way ANOVA, *p* = 0.549). C) Ladder walk task for *Pclo^wt/wt^*, 859 *Pclo^wt/gt^* and *Pclo^gt/gt^* rats for 3 trials over 1 day. *Pclo^gt/gt^* rats had a higher rate of stepping errors (ladder rung foot slips/misses) than *Pclo^wt/wt^* and *Pclo^wt/gt^* rats (forelimb error/step: *Pclo^wt/wt^* = 0.09 ± 0.00925, *Pclo^wt/gt^* = 0.148 ± 0.0263, *Pclo^gt/gt^* = 0.297 ± 0.0145, n = 4, one-way ANOVA, *p\*\** = 0.002, *p^#^ =* 0.0157 (where * **=** *p* value between *Pclo^wt/w^* and *Pclo^gt/gt^* and ^#^ = *p* value between *Pclo^wt/wt^* and *Pclo^wt/gt^*); hindlimb error/step: *Pclo^wt/wt^* = 0.0563 ± 0.0221, *Pclo^wt/gt^* = 0.0663 ± 0.00877, *Pclo^gt/gt^* = 0.238 ± 0.0543, n = 4; one-way ANOVA with Bonferroni correction, *p\** = 0.0135, *p^#^ =* 0.0186). D) Open field task for *Pclo^wt/wt^*, *Pclo^wt/gt^* and *Pclo^gt/gt^* rats for 1 trial each over 1 day. *Pclo^gt/gt^* rats performed fewer rearing behaviors than *Pclo^wt/wt^* and significantly less than *Pclo^wt/gt^* rats in the perimeter sectors of the arena (*Pclo^wt/wt^* = 64.25 ± 18.31 events; *Pclo^wt/gt^* = 76.25 ± 11.3 events; *Pclo^gt/gt^* = 22 ± 4.203 events; n = 4 rats per genotype, one-way ANOVA with Bonferroni correction, *p^#^* = 0.0427 (where ^#^ = *p* value between *Pclo^wt/wt^* and *Pclo^wt/gt^*). Other behaviors such as crossing the open field and grooming were not different between the three groups (crossing: *Pclo^wt/wt^* = 178.3 ± 29.68 events; *Pclo^wt/gt^* = 164.5 ± 45.51 events; *Pclo^gt/gt^* = 235.5 ± 27.11 events, n = 4 rats per genotype, one-way ANOVA with Bonferroni correction, *p* = 0.828; grooming: *Pclo^wt/wt^* = 2.75 ± 0.75 events; *Pclo^wt/gt^* = 2 ± 1.08 events; *Pclo^gt/gt^* = 5 ± 1.155 events, n = 4 rats per genotype, one-way ANOVA with Bonferroni correction, *p* = 0.488) E). Locomotor activity of *Pclo^wt/wt^*, *Pclo^wt/gt^* and *Pclo^gt/gt^* rats during the 12h dark phase. *Pclo^gt/gt^* rats covered a more than 50% longer distance than *Pclo^wt/wt^* and *Pclo^wt/gt^* rats (*Pclo^wt/wt^* = 135.7 ± 11.25, n = 10; *Pclo^wt/gt^* = 129.3 ± 10.44, n = 4; *Pclo^gt/gt^* = 217.8 ± 14.33, n = 12; one-way ANOVA, *p\*\*\** = 0.0002). Data points are individual means over 15 nights. F) Performance of *Pclo^wt/wt^*, *Pclo^wt/gt^* and *Pclo^gt/gt^* rats during the isometric pull-task (handle position 11 mm inside the cage). Only 4 out of 11 *Pclo^gt/gt^* rats succeeded at the 60 g force threshold and *Pclo^gt/gt^* rats pulled with a significantly lower success rate at all force thresholds as compared to *Pclo^wt/wt^* and *Pclo^wt/gt^*. (30g: *Pclo^wt/wt^* = 86.722 ± 2.3, n = 10; *Pclo^wt/gt^* = 84.374 ± 6.324, n=4; *Pclo^gt/gt^* = 61.044 ± 5.928, n = 11; 2 way ANOVA with Bonferroni correction, *p*\*\* = 0.0018; 40g: *Pclo^wt/wt^* = 83.603 ± 2.86, n = 10; *Pclo^wt/gt^* = 85.136 ± 7.373, n=4; *Pclo^gt/gt^* = 47.804 ± 6.897, n = 10; two-way ANOVA with Bonferroni correction, *p*\*\*\*\* < 0.0001, *p*^###^ = 0.0007; 50g: *Pclo^wt/wt^* = 80.49 ± 3.442, n = 10; *Pclo^wt/gt^* = 71.930 ± 8.695, n=4; *Pclo^gt/gt^* = 33.647 ± 6.802, n = 10; two-way ANOVA with Bonferroni correction, *p*\*\*\*\* < 0.0001, *p*^###^ = 0.0005 60g: *Pclo^wt/wt^* = 69.186 ± 4.99, n = 10; *Pclo^wt/gt^* = 70.409 ± 10.181, n=3; *Pclo^gt/gt^* = 30.003 ± 11.359, n = 4; 2 way ANOVA with Bonferroni correction, *p*\*\*\* = 0.0004, *p*^##^ = 0.0056). Error bars represent SEM. Data points represent individual rats.

In addition to deficits in motor ability, *Pclo^gt/gt^* rats displayed an increased frequency in front and rear foot stepping errors during ladder rung walking compared to *Pclo^wt/wt^* and *Pclo^wt/gt^* rats (Figure 9C). In open field tests, no significant differences were recorded between *Pclo^wt/wt^*, *Pclo^wt/gt^* and *Pclo^gt/gt^* rats for peripheral line crossing or self-grooming events (Figure 9D). However, *Pclo^gt/gt^* rats displayed 2- to 3-fold decreases in peripheral rearing events compared to *Pclo^wt/wt^* and *Pclo^wt/gt^* animals (Figure 9D). Deficits in *Pclo^gt/gt^* rat performance during rotarod, ladder rung and open field tests (Figure 9A-D) reflected recessive traits, consistent with dysfunction in proprioceptive sensation and motor control (Curzon et al., 2009).

Alongside traditional tests for motor coordination, we tested the behavior of *Pclo^wt/wt^*, *Pclo^wt/gt^* and *Pclo^gt/gt^* rats in a home cage setup, the OptiMan (Operator Independent Motor-analysis) system, where animals can be monitored without interference from experimenters. Rats were tagged with a radio frequency identification chip that allowed for tracking of locomotor activity while they were also required to complete an isometric pull task that requires precise and finely controlled movements. In the home cage, the *Pclo^gt/gt^* rats were more active and covered about twice the distance than *Pclo^wt/wt^* rats (Figure 9E) during each measurement. In the isometric pull task, performance of *Pclo^gt/gt^* rats was significantly lower than the performance of *Pclo^wt/wt^* rats, quantified using four different force thresholds (Figure 9F). Taken together, *Pclo^gt/gt^* rats show clear motor deficits, very similar to symptoms seen in PCH3 patients (Rajab et al., 2003).

## Discussion

Our study demonstrates that Piccolo LOF causes alterations in brain anatomy. In particular, the cerebrum, pons, brainstem and cerebellum are severely reduced in size whereas the ventricles are increased (Figure 1). These changes are associated with reductions in cerebellar and pontine cell numbers and perturbations in cerebellar CF and MF afferents (Figure 4 and 5). These changes are predicted to adversely affect cerebellar function - supported by changes in synaptic transmission and motor control in *Pclo^gt/gt^* rats. Interestingly, the changes in brain morphology resemble changes in children with PCH3, recently linked to a SNP in the PCLO gene (Ahmed et al., 2015). In addition to brain atrophy, children with PCH3 have, amongst other symptoms, cognitive and motor deficits as well as seizures. Given that seizures were also observed in *Pclo^gt/gt^* rats (Medrano et al., 2019), we postulate that *Pclo^gt/gt^* rats represent a model to study the underlying mechanisms of this devastating disease.

Similar to patients with PCH3, *Pclo^gt/gt^* rats have smaller cerebella, brainstem and pontine nuclei. In the cerebellum, a striking change was the thickness of the ML and CGL (Figure 3). The latter was associated with fewer total GCs, which could result in fewer PFs and a thinner ML. The orientation of PCs and the ramification of their dendritic arbors were largely unchanged. At present, it is unclear why there are fewer GCs in *Pclo^gt/gt^* cerebella. During normal development, several factors including Sonic hedgehog (Wallace, 1999; Miyashita et al., 2017) and Notch (Solecki et al., 2001) are known to control the proliferation of GCs. How Piccolo loss influences these and related signaling pathways is unclear; though Piccolo and Bassoon have been shown to regulate the activity-dependent translocation of c-terminal binding protein 1 (CTPB1) to the nucleus and thereby the expression of neuronal genes (Ivana et al., 2015). CTPB1 and CTPB2 are highly expressed in the cerebellum and their function and/or localization could be affected by Piccolo LOF (Hubler et al., 2012; Ivanova et al., 2015). There is a clear rationale for exploring this phenotype further.

In the ML, CF morphology and arrangement appear to be normal, though a higher innervation of PC dendrites by CFs can be observed (Figure 4). One possible explanation for this hyper-innervation is that homeostatic changes to the network, such as heterosynaptic competition with PFs for PC dendrite territory (Hashimoto et al., 2009; Ichikawa et al., 2016) could contribute to these differences in CF distribution and therefore the functionality of the cerebellum. This could affect normal pruning mechanisms of CF during development.

Our study also revealed that the pontine nuclei was dramatically decreased in size in *Pclo^gt/gt^* animals. This is note-worthy as afferent MFs from the pons and other brainstem nuclei are the primary excitatory input onto GCs, forming elaborate rosette synapses (Voogd and Glickstein, 1998). The smaller size of the pons suggests a net reduction in MF input into the cerebellum that appears to correspond to the reduced number of GCs (Figure 3). However, it is unclear how the loss of Piccolo could influence the number of neurons in the pons.

Surprisingly, our histological studies revealed that MF boutons across all lobes of the cerebellum were reduced in size (Figure 5). This finding was supported by our EM studies revealing that MF glomeruli are severely reduced in size, potentially resulting in smaller SV clusters (Figure 7). These observations suggest that the development/maturation of MFs from the pons and other brainstem nuclei are muted in *Pclo^gt/gt^* animals. Additionally, electrophysiological changes (Figure 8) indicate that the network has triggered compensatory changes to overcome a smaller size or impaired strength of MFs reaching the GC layer, for instance by increasing both input resistance and release probability (Turrigiano, 2012).

Our electron micrographs also showed what could be disturbed synaptic integrity in *Pclo^gt/gt^* MF boutons with more CCVs being present (Figure 7), suggesting that the loss of Piccolo may also affect SV recycling. These observations are in accordance with recent findings by Ackermann et al. (2019) of hippocampal synapses from *Pclo^gt/gt^* animals. Here, it was reported that the loss of Piccolo had a dramatic effect on the recycling of SV proteins through a functional block in the formation of early endosomes from endocytic vesicles, due to defects in the activation of Rab5 via a Piccolo-dependent loss of Pra1 from synapses. Although beyond the current study, it is likely that this endocytic defect in the recycling of SV proteins could also contribute to altered size function of MF synapses, especially given their high frequency transmission and therefore need for high SV turnover (Byczkowicz et al., 2018).

GCs also receive inhibitory modulation from Golgi cells (Eccles et al., 1966) and it is well appreciated that GABA_A_ receptors at the MF synapse contain α6 subunits (Nusser et al., 1996). Intriguingly, *Pclo^gt/gt^* rats display a reduced expression of the GABA_A_α6 subunit at MF rosettes (Figure 6). This is in line with findings from Medrano et al. (2019), who performed RNA sequencing on the *Pclo^gt/gt^* rats and found a severe reduction in GABA_A_α6 subunit gene expression.

At present, it is unclear why levels of these subunits are reduced. One possibility is that it reflects a homeostatic change within cerebellar circuitry to compensate for a reduced excitatory input from MFs (Figure 5). This concept is supported by electrophysiological data showing that the input resistance of GCs is higher as is mEPSC frequency and EPSC amplitudes (Figure 8). This condition might arise to compensate for the smaller MF terminals reaching the GCL, which could initially have weaker output properties. Alternatively, reduced GABA_A_α6 expression could be due to lower levels of BDNF, which is secreted from precerebellar neurons, at MF terminals (Chen et al., 2016) and is necessary to promote the formation of GABAergic synapses onto GCs. Though a role of Piccolo in the secretion of BDNF has not been investigated, the expression of Bassoon, which shares significant functional redundancy with Piccolo, is linked to presynaptic levels of BDNF (Heyden et al., 2011). In this regard, the remaining Bassoon protein could suppress BDNF secretion in MF terminals, altering the maturation of the glomeruli and GABA_A_α6 expression.

An important question not addressed by our studies is why MF boutons are smaller. By several measures mentioned above, it would appear that the MF glomeruli are less mature. Key regulators of MF maturation are members of the Wnt family, a group of target-derived factors which accelerate neuronal maturation or directly induce synapse formation (Scheiffele, 2003; Waites et al., 2005). Interestingly, mouse cerebellar MFs lacking Wnt7a show a similar reduction in MF size and complexity, as we observed in *Pclo^gt/gt^* (Hall et al., 2000). However, it is important to note that in the case of Wnt7a loss, MF synapse size catches up to WT size by age P15, which points to a lag in maturation and does not quite seem to be the case in our *Pclo^gt/gt^* rats. Knockout of Disheveled1 (Dvl1), a downstream target of Wnt (Salinas and Zou, 2008) also shows a reduction in MF cluster size, indicating that presynaptic Dvl1 is a necessary step in the Wnt signaling cascade, underscoring the importance of these proteins for MF-GC synapse formation (Ahmad-Annuar et al., 2006). Like Dvl1, Piccolo is located at the AZ of the presynaptic terminal and regulates F-actin assembly and synaptic transmission through its interaction with Daam1 and Profilin (Wagh et al., 2015). Daam1 is a formin and a known regulator/interaction partner of Dvl1 (Gao and Chen, 2010). It is thus possible that, in the absence of Piccolo, Dvl1 is not properly localized to presynaptic sites, preventing proper Wnt signaling and consequently causing the formation of smaller and less organized MF-GC synapses. Furthermore, analysis of *Pclo^gt/gt^* transcripts reveals that Wnt expression is reduced (Medrano et al., 2019). Thus, there appears to be at least two possible mechanisms that could contribute to smaller MF boutons in Piccolo knockout rats: defects in SV recycling and Wnt signaling. Clearly further studies are needed to explore these options.

A fundamental question raised by the anatomical, morphological and functional changes within the cerebellum in rats lacking Piccolo is whether these changes affect the functionality of the cerebellum. In behavioral tests, we observed that *Pclo^gt/gt^* rats performed significantly worse than both *Pclo^wt/wt^* and *Pclo^wt/gt^* littermates at motor function tasks, highlighting the recessive nature of the behavioral impairments. Our colleagues Medrano et al. (2019) also observed epileptic seizures and increased aggression in *Pclo^gt/gt^* rats, and a failure to reproduce due to impaired brain-gonad signaling.

Taken together, these data support the concept that Piccolo loss of function in patients with PCH3 could be causal for many of the observed phenotypes, including changes in the volume of brain structures and behavioral abnormalities such as impaired motor control and epileptic seizures (Rajab et al., 2003; Namavar et al., 2011). With regard to reduced cerebellar function, our studies highlight a prominent role for MF boutons which are not only smaller in size but with altered synaptic properties. Mechanistic studies, which probe how Piccolo loss contributes to these changes, should provide insights into the etiology of this devastating disease.

## Materials and Methods

### Generation of Piccolo KO rats (*Pclo^gt/gt^*)

Generation of mutant Piccolo rat strains: Mutant rat strains harbored Sleeping Beauty β-Geo trap transposons (Ivics et al., 2009), originally transmitted from a donor, recombinant rat spermatogonial stem cell library (Izsvak et al., 2010). Recipient males were bred with wildtype females to produce a random panel of mutant rat strains enriched with gene traps in protein coding genes (Izsvak et al., 2010). All experiments were approved by the Institutional Animal Care and Use Committee (IACUC) at UT-Southwestern Medical Center in Dallas, as certified by the Association for Assessment and Accreditation of Laboratory Animal Care International (AALAC) NIH OLAW Assurance # D16-00296.

### Characterization of pups and genotyping

All procedures for experiments involving animals, were approved by the animal welfare committee of Charité Medical University and the Berlin state government. P0-P2 rats were weighed using Kern 440-43N scales and measured for approximate length with a ruler. Genotyping of pups’ cortices later revealed their genetic identity.

P0-P2 pups were genotyped using a PCR based reaction. In brief, brain tissue was digested in lysis buffer (100 mM Tris-HCl (pH 8.0) with 10 mg/ml proteinase K, 100 mM NaCl) for 5 minutes at 55 °C, before inhibiting Proteinase K by incubation at 99 °C for 10 minutes. Samples were then centrifuged at 14,800 rpm for 2 minutes and 1 µl of supernatant was used for the PCR reaction as outlined below.

For determination of genotype for adult rats, earpieces were taken and digested overnight at 55 °C in SNET-buffer (400 mM NaCl, 1 % SDS, 200 mM Tris (pH 8.0), 5 mM EDTA) containing 10 mg/ml proteinase K. Proteinase K enzyme reaction was stopped incubating the samples for 10 min at 99 °C. The mixture was centrifuged for 2 min at 14,800 rpm. Supernatant was transferred into a fresh tube and DNA was precipitated by addition of 100 % isopropanol. Following samples were centrifuged for 15 min at 4 °C, 13,000 rpm. Precipitated DNA was washed once with 70 % ethanol and centrifuged again for 5 min at 13,000 rpm. Supernatant was discarded and the DNA pellet was air dried and resuspended in H2O. A PCR reaction with a specific primer combination was performed on isolated DNA. The following primers were used: Pclo KO F2: 3’ gcaggaacacaaaccaacaa 5’; Pclo KO R1: 3’ tgacctttagccggaactgt 5’; SBF2: 3’ tcatcaaggaaaccctggac 5’. The PCR reaction protocol was the following: 2 min 94 °C; 3 × (30 sec 94 °C, 60 °C 30 sec, 72 °C 30 sec); 35 × (94 °C 30 sec, 55 °C 30 sec, 72 °C 30 sec); 72 °C 10 min. Samples were mixed with a loading dye (New England BioLabs, MA, USA) and run on 2 % agarose gel (Serva, Heidelberg, Germany) at 110 V for 45 min. The gel was imaged using BioDocAnalyze UV transilluminator and BioDocAnalyze2.2 software.

### Western blot analysis

Brains from P0 – P2 pups were lysed in Lysis buffer (50 mM Tris-HCl, 150 mM NaCl, 5 mM EDTA, 1 % Triton X-100, 0.5 % Deoxycholate, protease inhibitor pH 7.5) and incubated on ice for 5 min. Samples were centrifuged at 13,000 rpm for 10 min at 4 °C. Afterwards the supernatant was transferred into a fresh tube and the protein concentration was determined using a BCA protein assay kit (Thermo Fisher scientific, Waltham, Massachusetts, USA). The same protein amounts for *Pclo^wt/wt^*, *Pclo^wt/gt^* and *Pclo^gt/gt^* samples were separated by SDS-PAGE and transferred onto nitrocellulose membranes (running buffer: 25 mM Tris, 190 mM Glycine, 0.1 % SDS, pH 8.3; transfer buffer: 25 mM Tris, 192 mM Glycine, 1 % SDS, 10 % Methanol for small proteins, 7 % Methanol for larger proteins pH 8.3). After the transfer, nitrocellulose membranes were blocked with 5 % milk in TBST (20 mM Tris pH 7.5, 150 mM NaCl, 0.1 % Tween 20) and incubated with primary antibodies in 3 % milk in TBST overnight at 4 °C. The following antibodies were used: Piccolo (1:1000; rabbit; Synaptic Systems, Göttingen, Germany; Cat# 142002), GABA_A_α6 receptor (1:300; Merck Darmstadt, Germany; Cat# 06-868) overnight. Nitrocellulose membranes were washed 3 times for 10 min with TBST and incubated with HRP-conjugated secondary antibodies for 1 h at RT (1:1000; Sigma-Aldrich, St. Louis, USA; Cat# NA9310 Cat# NA934). Membranes were washed 3 times for 10 min with TBST, afterwards secondary antibody binding was detected with ECL Western Blotting Detection Reagents (Thermo Fisher Scientific, Waltham, USA) and a Fusion FX7 image and analytics system (Vilber Lourmat, Collégien, France).

### Immunohistochemistry

Immunohistochemistry was performed on brain tissue from rats perfused with 4 % paraformaldehyde (Roth, Karlsruhe, Germany) dissolved in PB (80 mM Na2HPO4 (Roth, Karlsruhe, Germany), 20 mM NaH2PO4 (Bernd Kraft, Duisburg, Germany)) (PFA). Brains were extracted and placed in 4 % PFA overnight, cryoprotected in 15 % and then 30 % sucrose solution (Sigma-Aldrich, St. Louis, USA; dissolved in PB) for 24 hours each. Brains were snap-frozen by submersion in 2-methylbutane (Roth, Karlsruhe, Germany) cooled to −60 °C and then stored at −20 °C until use. Brains were cut para-sagittally or coronally with a Leica cryostat to either 20 µm thick sections and mounted on superfrost slides (Thermo Fisher Scientific, Waltham, USA), or 50 µm sections which were processed free-floating. Slides were left to dry for a minimum of one hour before storage at −20 °C and free-float sections were stored in antifreeze solution (30 % Ethylene glycol, 30 % Glycerol (Roth, Karlsruhe, Germany), 30 % ddH2O, 10 % 0.244M PO4 buffer (NaOH, NaH2PO4, Roth, Karlsruhe, Germany)).

Prior to staining at least 4 slides (each containing two sections) from each animal were left to equilibrate at room temperature (RT) for one hour. Sections were selected to encompass the range of the axis we were investigating. A hydrophobic barrier was created around sections using a DAKO pen (DAKO, Glostrup, Denmark) and sections were washed and permeabilized with TBS (20 mM Tris pH 7.5, 150 mM NaCl, Roth, Karlsruhe, Germany) with 0.025 % Triton X-100 (Roth, Karlsruhe, Germany) (TBST) for 3 × 5 min, prior to blocking with 10 % Normal Goat Serum (NGS, Sigma-Aldrich, St. Louis, USA) with 1 % Bovine Serum Albumin (BSA, Sigma-Aldrich, St. Louis, USA) in TBS. The following primary antibodies were used: Calbindin (1:750; rabbit; Synaptic Systems, Göttingen, Germany; Cat# 214002), GABA_A_ receptor α subunit (1:250; rabbit; Sigma-Aldrich, St. Louis, USA; Cat# G5544), MAP2 (1:1000; chicken; MilliporeSigma, Burlington, USA, Cat# AB5543), Piccolo (1:200; guinea pig; Synaptic Systems, Göttingen, Germany; Cat# 142 104), VGluT1 (1:1000; rabbit; Abcam, Cambridge, UK; Cat# ab104898), VGluT2 (1:250; guinea pig; Synaptic Systems, Göttingen, Germany; Cat# 13540419). VGluT2 (1:300; mouse monoclonal (epitope AA 566 to 582); Synaptic Systems, Göttingen, Germany; Cat# 135 421). Antibodies diluted in TBS with 1 % BSA were applied and left overnight at 4 °C. After 3 × 5 min washing with TBST, differently labeled secondary antibodies were used from Invitrogen (Thermo Fisher Scientific, Waltham, USA, dilution 1:1000), again diluted in 1 % BSA in TBS antibody solution and then applied for 1 hour at RT. Sections were then washed with TBS 2 × 10 min or, if desired, incubated in TBS with DAPI (Roth, Karlsruhe, Germany) for 30 min before washing again 2 × 10 min. Slides were coverslipped (24×50 mm coverslips, Menzel Gläser, Braunschweig, Germany) with Immu-Mount (Shandon-Thermo Scientific, Cheshire, UK) and sealed with clear nail polish once hardened.

### Nissl stain

Sections were washed 3 times in PBS then mounted onto superfrost slides and allowed to dry for 1-2 days. Slides were inserted into slide racks and passed through the following solutions: 95 % EtOH (Ethanol from Roth, Karlsruhe, Germany; diluted as appropriate with ddH2O) × 15 min, 70 % EtOH, 50 % EtOH, ddH2O, 10 min Blue counterstain (TACS from Trevigen, MD, USA), ddH2O and then dehydrated through 50 %, 70 % acid EtOH (1 % glacial acetic acid (Th. Geyer, Renningen, Germany) in 70 % EtOH), 95 % and 100 % EtOH before clearing in Roti Histol (Roth, Karlsruhe, Germany), coverslipped using Entellan mounting medium (Entellan, Darmstadt, Germany) and left to dry for 24 hours under a fume hood.

### Confocal z-stack image acquisition and processing

Images were acquired on a spinning disc confocal microscope (Zeiss Axio Observer.Z1 with Andor spinning disc and cobolt, omricron, i-beam laser) using a 40x (1.3 NA) and 100x (1.4 NA) Plan-Apochromat oil objective and an iXon ultra (Andor, Belfast, UK) camera controlled by iQ software (Andor, Belfast, UK). Sections for GABA_A_α6 analysis (Figure 7) were imaged with a Nikon Spinning Disk Confocal CSU-X using a 100x (1.45 NA) Plan Apo oil objective and an EMCCD camera with Andor Revolution SD System(Andor, Belfast, UK).

### Tile scan overview images

An upright microscope (Olympus BX61) was used to image fluorescently stained cerebellar sections. A CCD color camera was used with a 10x or 5x lens for overview pictures. CellSens software (Olympus, Hamburg, Germany) stitched multiple images together to give an overview of the whole cerebellar region.

### Image analysis

For image processing and analysis ImageJ/FIJI software was used (Schindelin et al., 2012). For analysis of *Pclo^wt/wt^* and *Pclo^gt/gt^* tissue sections were selected from the vermis and approximately every 10th slide laterally (each slide containing 2 sections) was analyzed. The best quality section per slide was imaged. For layer thickness, all lobes were measured and for closer analysis of MF size, CF coverage and GC density, images were taken from lobes I, III, V, VII and IX where possible. For GABA_a_ɑ6 subunit expression, 6 images were taken per slide, 4 slides per animal. Average signal intensity was measured for the whole field of view in the GABA_a_ɑ6 antibody channel and then normalized to the MAP2 channel for each image.

Layer ‘thickness’ was calculated per lobe by dividing the area of the layer by the inner length of the layer for both GCL and ML. For Purkinje cell density, the number of PCs per lobe was counted and divided by the length of the PC layer for that lobe. Data points represent ‘thickness’ values from individual lobes.

Granule cell density was calculated from 100x magnification single plane images stained with DAPI. GCs were counted per image by an experimenter blinded to the genotype of each image, and this number divided by the area of each image. Data points represent each image.

Mossy fiber cluster size was measured using a script within ImageJ to analyze particles stained with VGluT1 and/or VGluT2. The area of each puncta was measured and an average taken per image. Data points represent average cluster size of each image.

Climbing fiber innervation was assessed using VGluT2 and Calbindin staining. The molecular layer area was defined in the Calbindin channel, consisting of Calbindin-positive PC dendrites, and the VGluT2 channel was selected and pasted into a new imageJ file. The same script as for MF analysis was run to calculate the total area stained with VGluT2, divided by the area of the ML and x100 to give percentage coverage. Data presented represent percentage per image.

### Electron microscopy

*Pclo^wt/wt^* and *Pclo^gt/gt^* rats were anesthetized deeply with 33 mg/ml Ketamine (Inresa Arzneimittel GmbH, Freiburg, Germany) 830 µg/ml Xylavet (cp-pharma, Burgdorf, Germany) and perfused transcardially with 37 °C physiological saline for 3 to 4 min followed by a 0.1 M phosphate buffer containing 4 % paraformaldehyde and 0.05 % glutaraldehyde. Brains were stored in fixative overnight and subsequently sliced sagittally (350 µm) on a vibratome. Regions of interest were cut into small pieces and post-fixed in 1 % OsO4 and 0.1 M sodium cacodylate and en-bloc stained in 1 % uranyl acetate aqueous solution. Finally, samples were dehydrated and embedded in epoxy resin (Epon 812; EMS). Ultrathin sections were cut using an Ultracut ultramicrotome (UCT, Leica, Wetzlar, Germany) equipped with a diamond knife (Ultra 45, DiATOME, Hatfield, USA) and collected on formvar-coated 200-mesh copper grids (EMS). Sections were imaged in a Zeiss EM-900 Transmission Electron Microscope (Zeiss) operated at 80 keV and equipped with a 2K x 2K CCD camera. Data were analyzed blindly using the ImageJ software. Data points represent analysis from individual images obtained from 3 rats per genotype. For histological and EM data, a normality test was performed (D’Agostino-Pearson omnibus normality test). If successful then a student’s t-test was used to compare *Pclo^wt/wt^* to *Pclo^gt/gt^* rats, or alternatively a test for non-normal data (Mann-Whitney U test) was used.

### Electrophysiology methods

Cerebellar slices were prepared from 3-to-4 month-old *Pclo^wt/wt^* and *Pclo^gt/gt^* rats of either sex. Animals were treated in accordance with the German Protection of Animals Act and with the guidelines for the welfare of experimental animals issued by the European Communities Council; local authorities approved the experiments. Animals were anesthetized with isoflurane (Baxter, Deerfield, IL) followed by a rapid decapitation with a custom-built guillotine. The cerebellar vermis was quickly removed and mounted in a chamber filled with chilled extracellular solution. Parasagittal 300 µm slices were cut using a Leica VT1200 microtome (Leica Microsystems, Wetzlar, Germany), transferred to an incubation chamber at ~35 °C for 30 min and subsequently stored at RT. Artificial cerebrospinal fluid (ACSF) was used for slice cutting, storage, and experiments. ACSF contained: 125 mM NaCl, 25 mM NaHCO3, 2.5 mM KCl, 1.25 mM NaH2PO4, 1.1 mM CaCl2, 1 mM MgCl2, 3 mM Glucose, 17 mM Sucrose (~310 mOsm, pH 7.3 when bubbled with Carbogen (5% O2/95% CO2)). Patch pipettes were pulled from borosilicate glass (Science Products, Hofheim, Germany) using a DMZ Puller (Zeitz-Instruments, Martinsried, Germany). Patch pipettes had open-tip resistances of 6–9 MΩ. The intracellular solution contained: 150 mM K-gluconate, 10 mM NaCl, 10 mM K-HEPES, 3 mM Mg-ATP, 0.3 mM Na-GTP (300–305 mOsm, pH adjusted to 7.3 with KOH). In some of the experiments, the intracellular solution contained 10 µM of the fluorescence dye Atto594. Experiments were performed at 35–37 °C and slices were continuously superfused with ACSF containing 20 µM SR95531 and 40 µM D-(2R)-amino-5-phosphonovaleric acid (D-APV) to block Golgi-cell inhibition and NMDA-receptors, respectively. Atto594 was obtained from Atto-Tec (Atto-Tec, Siegen, Germany); all other chemicals were purchased from Sigma-Aldrich (St. Louis, MO).

### Current clamp recordings

To determine the input resistance, subthreshold current pulses were applied from −20 to +20 pA in 2pA steps. Action potentials were evoked in current-clamp mode by current pulses (amplitude 20-400 pA, duration 300 ms). The resistance of the solution-filled patch-pipettes was 24.9 ± 1 MΩ and 23 ± 1 MΩ for *Pclo^wt/wt^* and *Pclo^gt/gt^* rats respectively. Patch-clamp recordings were made using a HEKA EPC10/2 USB amplifier (HEKA Elektronik, Lambrecht/Pfalz, Germany). Data were sampled at 200 kHz. Measurements were corrected for a liquid junction potential of +16 mV.

#### Excitatory postsynaptic currents (EPSCs)

To measure evoked EPSCs, GCs were held at a holding potential of – 80 mV and Mossy fiber axons were stimulated at 1 Hz. The stimulation voltage ranged between 16 to 40 V for control and *Pclo^gt/gt^* animals. For spontaneous EPSCs GCs were held at −80 mV for around 3 minutes. Single events were detected using the Igor Pro extension NeuroMatic (Rothman and Silver, 2018b) tool for Event detection.

The current clamp data were analyzed using custom-made procedures in Igor Pro software (WaveMetrics, Oregon, USA) as described previously (PMID:31379501). In short, properties of action potentials of GCs were determined from the injected currents, that elicited the largest number of action potentials (APs). The action potential threshold was defined as the membrane voltage at which the first derivative exceeded 100 V s^−1^, the minimal AP peak was set as −20 mV and the minimal amplitude to 20 mV. All APs with a half-width smaller than 50 µs and higher than 500 µs were excluded. AP frequency and AP half-width were calculated from the first three APs. Membrane capacitance, resting membrane potential and series resistance were read from the amplifier software (HEKA) after achieving the whole-cell configuration. Input resistance (R_in_) was analyzed by plotting the steady-state voltage elicited by the subthreshold current injections against the injected current and a spline interpolation was performed to obtain the slope at the holding membrane potential (0 pA current injection).

#### Excitatory postsynaptic currents

To measure evoked EPSCs, GCs were held at a holding potential of – 80 mV and mossy fiber axons were stimulated extracellular with a second patch pipette at 1 Hz. For spontaneous EPSCs GCs, were held at −80 mV for around 3 minutes. Single events were detected using the template detection algorithm of the Igor Pro extension NeuroMatic (Rothman and Silver, 2018a).

EPSCs were analyzed with the Igor Pro software. The amplitude and the kinetics were determined from the average of 25 individual single EPSCs. To obtain the decay kinetics, single EPSCs were fitted with one or two exponentials. The weighted time constant was calculated as:

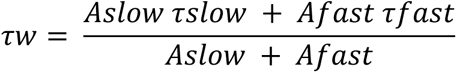

### Behavioral assessment

#### Rotarod task

The apparatus (IITC Life Science, Woodland Hills, CA) consisted of 5 semi-enclosed lanes and an elevated metal rod (9.525 cm diameter, 29.21 cm elevation) with a fine textured finish to enhance grip. For each trial, all rats were placed on the unmoving rod, allowed to stabilize their posture, and then rod rotation was initiated. Test parameters were: rotation direction, toward investigator to encourage rats to face away while walking; start speed, 4 rpm; top speed, 44 rpm; acceleration rate, 0.2 rpm/s (200 sec from start to top speed); max test duration, 300 s. Each rat’s trial ended when it fell from the rod, triggering the fall-detection sensor. Data was automatically recorded to a computer. Rats underwent 4 trials/day, with an inter-trial interval of at least 10 min for 4 consecutive days.

### Ladder rung task

Ladder rung tests were performed on cohorts of *Pclo^wt/wt^*, Pclo^wt/gt^ and *Pclo^gt/gt^* rats by methods previously described (Metz and Whishaw, 2009). Rats were trained to cross a plexiglass tunnel about 1 m long with metal rods provided at regular intervals as steps. Each step that the rat took was scored on the basis of paw placement on a 7 category scale with 0 being paw totally missing the rung and 6 being correct paw placement. The average score per pair of limbs was used to quantify step score. During the trial, the error per step was also quantified by number of low scoring steps (0-2) divided by the total number of steps that the rat took during the test with each pair of limbs (student’s t-test). Data were derived from 3 trials with 3 cm rung separation conducted on the same test day with at least a 10 min inter-trial interval, and group means were compared using one-way ANOVA and Bonferroni post-hoc tests.

### Grip test

To assess grip strength, rats were allowed to cling on to a support by forelimbs or hindlimbs and pulled (Curzon et al., 2009). The support is attached to a transducer that can measure the pull force being applied on the rat by the tester. During each trial, force that was necessary to be applied by the tester to release the grip of the rat was recorded. Three trials per limb pair were done and the means compared by student’s t-test. To assess grip strength, the rat’s torso was supported ventrally while both forelimbs were allowed to grasp a metal support bar, then the rat was pulled in a horizontal plane until the bar was released. Peak force was measured by a transducer attached to the support bar (San Diego Instruments, San Diego, USA). Nine trials per rat were conducted over two days (inter-trial interval at least 5 min) and group means were compared using one-way ANOVA and Bonferroni post-hoc tests.

### Open field test

The open field test (Curzon et al., 2009) was performed for each rat using a rectangular arena (91.44 cm × 60.96 cm) divided into 9 equal sectors (1 center, 8 perimeter). At the start of the experiment, the rat was placed in the center sector and allowed to explore for 10 min. The test was recorded by digital video for later analysis. One test was conducted per rat. The behaviors counted were crossing, rearing and grooming. Each crossing event was counted when all four limbs of the test subject crossed the boundary of one sector into another. Rearing was defined as the number of times the subject stood up on its hind limbs. Grooming was defined as the number of times a rat licked or scratched itself while remaining stationary. Events that occurred in the center versus perimeter sectors were tallied separately, and group means were analyzed for each region by one-way ANOVA and Bonferroni post-hoc tests.

### OptiMan system

Experiments were approved by the animal welfare committee of Charité Medical University and the Berlin state government. Four groups (22 male & 4 female) of *Pclo^wt/wt^*, Pclo^wt/gt^ and *Pclo^gt/gt^* rats were used in this study. Rats carried subcutaneous radio-frequency identification (RFID) tags in ventral location. Animals were group housed with 6 - 8 rats per group. Initially, all rats were habituated to the OptiMan multi-cage environment with open sorter gates for voluntary exploration. Then, automated sorting was activated so that only one rat could enter the operant chamber at a time. During pull-task training, force thresholds and handle positions were adapted every day for each session to the current skill level reached by a rat. Each session had 20 trials, and two to three individual sessions were given per day with an intersession interval of 30 to 60 min. Within a session, the maximum time interval a rat was allowed to remain inactive between trials was 6 - 8 min. A session ended when such inactivity occurred. The OptiMan (Operator Independent Motor-Analysis, PhenoSys) system consists of two interconnected group home cages (EU Type IV cage and 2000P) resting on an RFID sensor array plate that automatically tracks RFID tag movement within the cage, and thus the locomotor activity of individual rats, day and night. One of the home cages was connected via an electronic guillotine gate to a second cage resting on a balance that automatically determined the body mass of a rat when inside this cage. Individual animals voluntarily and self-motivated decided to visit the balance cage. Subsequent to the balance cage, and again separated by an electronic guillotine gate, was a cage compartment containing a horizontal ladder (1 m) with electronically monitored dual rungs on the left and right side. From this ladder compartment, an animal entered a cage containing the isometric pull-task. From there back to the home cage a rat passed over a force grid sensor array that sensed ground forces exerted by the paws.

### Isometric pull task

Rats were trained to pull a handle attached to a stationary force transducer with a predetermined force threshold upon which they received a sugar pellet reward. Upon a rat’s entry into the isometric pull task cage a sugar pellet was delivered into the reward tray. A session started when the rat retrieved this first reward, which led to the automatic closing of the entry gate, and to the motorized slide-in appearance of the force handle to its predefined position. During each session, 5 different handle positions were presented to a rat. These positions varied from 11 mm inside the cage wall (easiest position), to 7 mm outside of the cage wall (most difficult position). Handle positions changed automatically during a session through a motorized slide. The difficulty level within each handle position increased stepwise by increasing the force threshold for pellet release. This started at 30 g pull-force and was increased to 40 g, 50 g and finally 60 g. A trial started when a pull-force of 5 g was sensed. From then on, the pull-force was sampled for a duration of 4 sec. If a rat reached the force threshold within a 2 sec time interval then a trial was successful and a reward delivered. The schedule then advanced to the next level of difficulty by either increasing force threshold or moving the handle one position further towards the outside. Thus, a rat needed a minimum of 20 trials to complete a session with 5 different handle positions and 4 different force thresholds at each position. If the threshold was not reached then a trial was considered unsuccessful and the rat had to continue with its next trial with unchanged conditions. Experiments lasted for 15 experimental days with 2 - 3 daily sessions per individual.

### Animal welfare

All animals were treated and cared for in accordance with national and institutional guidelines:

Generation of mutant Piccolo rat strains and behavioral experiments Figure 10 a-e: Rat protocols were approved by the Institutional Animal Care and Use Committee (IACUC) at UT-Southwestern Medical Center in Dallas, as certified by the Association for Assessment and Accreditation of Laboratory Animal Care International (AALAC), permit number: NIH OLAW Assurance # D16-00296.

Western blotting, immunohistochemistry, electron microscopy and behavioral experiments with Optiman setup (Figure 10 d and e): Animals were treated in accordance with the German Protection of Animals Act (TierSchG §4 Abs. 3); all procedures for experiments involving animals were approved by the animal welfare committee of Charité Medical University and the Berlin state government, permit number: T 0036/14.

Electrophysiology: Animals were treated in accordance with the German Protection of Animals Act (TierSchG §4 Abs. 3) and with the guidelines for the welfare of experimental animals issued by the European Communities Council Directive of 24. November 1986 (86/609/EEC). The local authorities approved the experiments (Landesdirektion Leipzig), permit number: T24/18

### Experimental design and statistical analysis

GraphPad Prism (RRID:SCR_002798) was used to analyze and represent data. Statistical design, sample sizes and tests for each experiment can be found in the figure legends.

## Acknowledgments

We would like to thank Susanne Wegmann and Eckart Gundelfinger for discussion and valuable comments on the manuscript; Anny Kretschmer and Katja Czieselsky for technical assistance. The work was supported by German Center for Neurodegenerative Diseases (DZNE), the Federal Government of Germany (DFG) SFB958 to CCG and (DFG) EXC 257 for the Center of Excellence NeuroCure to YW. Work to generate *Pclo^gt/gt^* rats was supported by National institute of Health R24RR03232601 & R24OD011108 to FKH. Neurological analyses on Piccolo mutant rats were conducted by The Neuro-Models Facility (EJP, LBG) at UT Southwestern Medical Center, and supported by the Haggerty Center for Brain Injury and Repair.

## Notes

**Conflict of interest statement:** The authors declare no competing financial interests.

## References

Ackermann F, Schink KO, Bruns C, Izsvak Z, Hamra FK, Rosenmund C, Garner CC (2019) Critical role for Piccolo in synaptic vesicle retrieval. Elife 8.

Ahmad-Annuar A, Ciani L, Simeonidis I, Herreros J, Fredj NB, Rosso SB, Hall A, Brickley S, Salinas PC (2006) Signaling across the synapse: a role for Wnt and Dishevelled in presynaptic assembly and neurotransmitter release. J Cell Biol 174:127–139.

Ahmed MY, Chioza BA, Rajab A, Schmitz-Abe K, Al-Khayat A, Al-Turki S, Baple EL, Patton MA, Al-Memar AY, Hurles ME, Partlow JN, Hill RS, Evrony GD, Servattalab S, Markianos K, Walsh CA, Crosby AH, Mochida GH (2015) Loss of PCLO function underlies pontocerebellar hypoplasia type III. Neurology 84:1745–1750.

Apps R, Garwicz M (2005) Anatomical and physiological foundations of cerebellar information processing. Nat Rev Neurosci 6:297–311.

Brickley SG, Cull-Candy SG, Farrant M (1996) Development of a tonic form of synaptic inhibition in rat cerebellar granule cells resulting from persistent activation of GABAA receptors. J Physiol 497 (Pt 3):753–759.

Byczkowicz N, Ritzau-Jost A, Delvendahl I, Hallermann S (2018) How to maintain active zone integrity during high-frequency transmission. Neurosci Res 127:61–69.

Cases-Langhoff C, Voss B, Garner AM, Appeltauer U, Takei K, Kindler S, Veh RW, De Camilli P, Gundelfinger ED, Garner CC (1996) Piccolo, a novel 420 kDa protein associated with the presynaptic cytomatrix. Eur J Cell Biol 69:214–223.

Chen AI, Zang K, Masliah E, Reichardt LF (2016) Glutamatergic axon-derived BDNF controls GABAergic synaptic differentiation in the cerebellum. Sci Rep 6:20201.

Curzon P, Zhang M, Radek RJ, Fox GB (2009) The Behavioral Assessment of Sensorimotor Processes in the Mouse: Acoustic Startle, Sensory Gating, Locomotor Activity, Rotarod, and Beam Walking. In: Methods of Behavior Analysis in Neuroscience (nd, Buccafusco JJ, eds). Boca Raton (FL).

Durmaz B, Wollnik B, Cogulu O, Li Y, Tekgul H, Hazan F, Ozkinay F (2009) Pontocerebellar hypoplasia type III (CLAM): Extended phenotype and novel molecular findings. J Neurol 256:416–419.

Eccles JC, Llinás R, Sasaki K (1966) The mossy fibre-granule cell relay of the cerebellum and its inhibitory control by Golgi cells. Experimental Brain Research 1:82–101.

Fenster SD, Garner CC (2002) Gene structure and genetic localization of the PCLO gene encoding the presynaptic active zone protein Piccolo. Int J Dev Neurosci 20:161–171.

Fenster SD, Chung WJ, Zhai R, Cases-Langhoff C, Voss B, Garner AM, Kaempf U, Kindler S, Gundelfinger ED, Garner CC (2000) Piccolo, a presynaptic zinc finger protein structurally related to bassoon. Neuron 25:203–214.

Gao C, Chen YG (2010) Dishevelled: The hub of Wnt signaling. Cell Signal 22:717–727.

Gundelfinger ED, Reissner C, Garner CC (2015) Role of Bassoon and Piccolo in Assembly and Molecular Organization of the Active Zone. Front Synaptic Neurosci 7:19.

Hall AC, Lucas FR, Salinas PC (2000) Axonal remodeling and synaptic differentiation in the cerebellum is regulated by WNT-7a signaling. Cell 100:525–535.

Hashimoto K, Kano M (1998) Presynaptic origin of paired-pulse depression at climbing fibre-Purkinje cell synapses in the rat cerebellum. J Physiol 506 (Pt 2):391–405.

Hashimoto K, Ichikawa R, Kitamura K, Watanabe M, Kano M (2009) Translocation of a “winner” climbing fiber to the Purkinje cell dendrite and subsequent elimination of “losers” from the soma in developing cerebellum. Neuron 63:106–118.

Heyden A, Ionescu MCS, Romorini S, Kracht B, Ghiglieri V, Calabresi P, Seidenbecher C, Angenstein F, Gundelfinger ED (2011) Hippocampal enlargement in Bassoon-mutant mice is associated with enhanced neurogenesis, reduced apoptosis, and abnormal BDNF levels. Cell Tissue Res 346:11–26.

Homanics GE, Ferguson C, Quinlan JJ, Daggett J, Snyder K, Lagenaur C, Mi Z-P, Wang X-H, Grayson DR, Firestone LL (1997) Gene Knockout of the α6 Subunit of the γ-Aminobutyric Acid Type A Receptor: Lack of Effect on Responses to Ethanol, Pentobarbital, and General Anesthetics. Molecular Pharmacology 51:588–596.

Hubler D, Rankovic M, Richter K, Lazarevic V, Altrock WD, Fischer KD, Gundelfinger ED, Fejtova A (2012) Differential spatial expression and subcellular localization of CtBP family members in rodent brain. PLoS One 7:e39710.

Human Protein Atlas (2015) PCLO. In.

Ichikawa R, Hashimoto K, Miyazaki T, Uchigashima M, Yamasaki M, Aiba A, Kano M, Watanabe M (2016) Territories of heterologous inputs onto Purkinje cell dendrites are segregated by mGluR1-dependent parallel fiber synapse elimination. Proc Natl Acad Sci U S A 113:2282–2287.

Ivanova D, Dirks A, Montenegro-Venegas C, Schone C, Altrock WD, Marini C, Frischknecht R, Schanze D, Zenker M, Gundelfinger ED, Fejtova A (2015) Synaptic activity controls localization and function of CtBP1 via binding to Bassoon and Piccolo. EMBO J 34:1056–1077.

Ivics Z, Li MA, Mates L, Boeke JD, Nagy A, Bradley A, Izsvak Z (2009) Transposon-mediated genome manipulation in vertebrates (vol 6, pg 415, 2009). Nat Methods 6:546–546.

Izsvak Z, Frohlich J, Grabundzija I, Shirley JR, Powell HM, Chapman KM, Ivics Z, Hamra FK (2010) Generating knockout rats by transposon mutagenesis in spermatogonial stem cells. Nat Methods 7:443–445.

Jakab RL, Hamori J (1988) Quantitative Morphology and Synaptology of Cerebellar Glomeruli in the Rat. Anat Embryol 179:81–88.

Leto K et al. (2016) Consensus Paper: Cerebellar Development. Cerebellum 15:789–828.

Maex R, De Schutter E (1998) Synchronization of golgi and granule cell firing in a detailed network model of the cerebellar granule cell layer. J Neurophysiol 80:2521–2537.

Maricich SM, Aqeeb KA, Moayedi Y, Mathes EL, Patel MS, Chitayat D, Lyon G, Leroy JG, Zoghbi HY (2011) Pontocerebellar Hypoplasia: Review of Classification and Genetics, and Exclusion of Several Genes Known to Be Important for Cerebellar Development. Journal of Child Neurology 26:288–294.

Medrano GA, Singh M, Plautz EJ, Good LB, Chapman KM, Chaudhary J, Jaichander P, Powell HM, Pudasaini A, Shelton JM, Richardson JA, Xie X-J, Ivics Z, Braun C, Ackermann F, Garner CC, Izsvák Z, Hamra FK (2019) Mutant screen reveals depression-associated *Piccolo’s* control over brain-gonad cross talk and reproductive behavior. bioRxiv:405985.

Metz GA, Whishaw IQ (2009) The ladder rung walking task: a scoring system and its practical application. J Vis Exp.

Miyashita S, Adachi T, Yamashita M, Sota T, Hoshino M (2017) Dynamics of the cell division orientation of granule cell precursors during cerebellar development. Mech Dev 147:1–7.

Miyazaki T, Fukaya M, Shimizu H, Watanabe M (2003) Subtype switching of vesicular glutamate transporters at parallel fibre-Purkinje cell synapses in developing mouse cerebellum. Eur J Neurosci 17:2563–2572.

Muller TM, Gierke K, Joachimsthaler A, Sticht H, Izsvak Z, Hamra FK, Fejtova A, Ackermann F, Garner CC, Kremers J, Brandstatter JH, Regus-Leidig H (2019) A Multiple Piccolino-RIBEYE Interaction Supports Plate-Shaped Synaptic Ribbons in Retinal Neurons. J Neurosci 39:2606–2619.

Namavar Y et al. (2011) Clinical, neuroradiological and genetic findings in pontocerebellar hypoplasia. Brain 134:143–156.

Nusser Z, Sieghart W, Somogyi P (1998) Segregation of different GABAA receptors to synaptic and extrasynaptic membranes of cerebellar granule cells. J Neurosci 18:1693–1703.

Nusser Z, Sieghart W, Stephenson F, Somogyi P (1996) The alpha 6 subunit of the GABAA receptor is concentrated in both inhibitory and excitatory synapses on cerebellar granule cells. The Journal of Neuroscience 16:103–114.

Rajab A, Mochida GH, Hill A, Ganesh V, Bodell A, Riaz A, Grant PE, Shugart YY, Walsh CA (2003) A novel form of pontocerebellar hypoplasia maps to chromosome 7q11-21. Neurology 60:1664–1667.

Regus-Leidig H, Fuchs M, Lohner M, Leist SR, Leal-Ortiz S, Chiodo VA, Hauswirth WW, Garner CC, Brandstatter JH (2014) In vivo knockdown of Piccolino disrupts presynaptic ribbon morphology in mouse photoreceptor synapses. Front Cell Neurosci 8:259.

Rothman JS, Silver RA (2018a) NeuroMatic: An Integrated Open-Source Software Toolkit for Acquisition, Analysis and Simulation of Electrophysiological Data. Front Neuroinform 12:14.

Rothman JS, Silver RA (2018b) NeuroMatic: An Integrated Open-Source Software Toolkit for Acquisition, Analysis and Simulation of Electrophysiological Data. Front Neuroinform 12.

Rudnik-Schoneborn S, Barth PG, Zerres K (2014) Pontocerebellar hypoplasia. Am J Med Genet C Semin Med Genet 166C:173–183.

Salinas PC, Zou YM (2008) Wnt signaling in neural circuit assembly. Annual Review of Neuroscience 31:339–358.

Scheiffele P (2003) Cell-cell signaling during synapse formation in the CNS. Annual Review of Neuroscience 26:485–508.

Schindelin J, Arganda-Carreras I, Frise E, Kaynig V, Longair M, Pietzsch T, Preibisch S, Rueden C, Saalfeld S, Schmid B, Tinevez JY, White DJ, Hartenstein V, Eliceiri K, Tomancak P, Cardona A (2012) Fiji: an open-source platform for biological-image analysis. Nat Methods 9:676–682.

Shinoda Y, Sugiuchi Y, Futami T, Izawa R (1992) Axon Collaterals of Mossy Fibers from the Pontine Nucleus in the Cerebellar Dentate Nucleus. Journal of Neurophysiology 67:547–560.

Sillitoe R, Fu, Y., Watson, C. (2012) Chapter 11 - Cerebellum: Academic Press.

Solecki DJ, Liu XL, Tomoda T, Fang Y, Hatten ME (2001) Activated Notch2 signaling inhibits differentiation of cerebellar granule neuron precursors by maintaining proliferation. Neuron 31:557–568.

Turrigiano G (2012) Homeostatic synaptic plasticity: local and global mechanisms for stabilizing neuronal function. Cold Spring Harb Perspect Biol 4:a005736.

Voogd J, Glickstein M (1998) The anatomy of the cerebellum. Trends Neurosci 21:370–375.

Wagh D, Terry-Lorenzo R, Waites CL, Leal-Ortiz SA, Maas C, Reimer RJ, Garner CC (2015) Piccolo Directs Activity Dependent F-Actin Assembly from Presynaptic Active Zones via Daam1. PLoS One 10:e0120093.

Waites CL, Craig AM, Garner CC (2005) Mechanisms of vertebrate synaptogenesis. Annual Review of Neuroscience 28:251–274.

Waites CL, Leal-Ortiz SA, Okerlund N, Dalke H, Fejtova A, Altrock WD, Gundelfinger ED, Garner CC (2013) Bassoon and Piccolo maintain synapse integrity by regulating protein ubiquitination and degradation. EMBO J 32:954–969.

Wallace VA (1999) Purkinje-cell-derived Sonic hedgehog regulates granule neuron precursor cell proliferation in the developing mouse cerebellum. Curr Biol 9:445–448.

Xu-Friedman MA, Regehr WG (2003) Ultrastructural contributions to desensitization at cerebellar mossy fiber to granule cell synapses. Journal of Neuroscience 23:2182–2192.

Zelnik N, Dobyns WB, Forem SL, Kolodny EH (1996) Congenital pontocerebellar atrophy in three patients: clinical, radiologic and etiologic considerations. Neuroradiology 38:684–687.

Zhai RG, Vardinon-Friedman H, Cases-Langhoff C, Becker B, Gundelfinger ED, Ziv NE, Garner CC (2001) Assembling the presynaptic active zone: a characterization of an active one precursor vesicle. Neuron 29:131–143.

